# How far to build it before they come? Analyzing the use of the Field of Dreams hypothesis to bull kelp restoration

**DOI:** 10.1101/2021.10.27.466118

**Authors:** Jorge Arroyo-Esquivel, Marissa L. Baskett, Meredith McPherson, Alan Hastings

## Abstract

In restoration ecology, the Field of Dreams Hypothesis posits that restoration efforts that create a suitable environment could lead to eventual recovery of the remaining aspects of the ecosystem through natural processes. Natural processes following partial restoration has lead to ecosystem recovery in both terrestrial and aquatic systems. However, understanding the efficacy of a “field of dreams” approach requires comparison of different approaches to partial restoration in terms of spatial, temporal, and ecological scale to what would happen with more comprehensive restoration efforts. We explore the relative effect of partial restoration and ongoing recovery on restoration efficacy with a dynamical model based on temperate rocky reefs in Northern California. We analyze our model for both the ability and rate of bull kelp forest recovery under different restoration strategies. We compare the efficacy of a partial restoration approach with a more comprehensive restoration effort by exploring how kelp recovery likelihood and rate change with varying intensities of urchin removal and kelp outplanting over different time periods and spatial scales. We find that, for the case of bull kelp forests, setting more favorable initial conditions for kelp recovery through implementing both urchin harvesting and kelp outplanting at the start of the restoration project has a bigger impact on the kelp recovery rate than applying restoration efforts through a longer period of time. Therefore partial restoration efforts, in terms of spatial and temporal scale, can be significantly more effective when applied across multiple ecological scales in terms of both the capacity and rate of achieving the target outcomes.

## 1 Introduction

One of the main challenges in restoration ecology is understanding the intensity and extent of the efforts required to achieve restoration goals (Bradshaw, 1996). The idea that partial restoration might be effetive is embodied in the Field of Dreams hypothesis (Palmer et al., 1997), which postulates that setting up favorable conditions for restoration at the beginning of the project can be enough to promote the natural processes that will lead to a successful restoration effort. Partial restoration has been successful in cases such as short-term habitat enhancement through a one-time coral reef transplantation, which then enhanced a longer-term natural recovery of coral (Maya et al., 2016), or reintroduction of former native species in degraded systems, which leads to increases in species richness in the community (Richardson et al., 2010). However, partial restoration may not always be effective at achieving management goals. In some cases the resulting community may not be desirable due to a lower diversity than the target community (Wodika and Baer, 2015), or stochasticity may bring similar ecosystems to completely different states, making further restoration efforts necessary if one of the states is undesirable (Trowbridge, 2007).

These examples raise the question of under what conditions engaging in partial restoration efforts and then relying on natural processes for ecosystem recovery can achieve restoration goals. This question can be explored in terms of partial restoration efforts occurring at three different scales (Wiens, 1992; Acosta et al., 2018). First, considering the spatial scale, partial restoration depends on the extent of the restoration effort that will then lead to natural recovery of the rest of the region through spread. Second, considering the temporal scale, partial restoration arises from performing restoration efforts at a short-term timeframe and longer term recovery following from natural dynamics. Third, considering the ecological scale, partial restoration arises from targeting a species or component of the community (e.g. reintroducing a foundational or early successional species, or removing a pest species), and then recovery of additional species in the community occurs naturally (e.g. through succession). Existing evaluations of partial restoration have explored different scales. For example, Stoddard et al. (2019) tested the ecological scale of restoration in terms of whether restoring dune vegetation could lead to the natural recovery of beach mice. They found that beach mice occupied restored habitats almost as frequently as natural habitats. In addition, a meta-analysis by Katwijk et al. (2016) of seagrass restoration found that as the spatial scale increases, the likelihood of restoration success increases as well. Therefore, different aspects of partial restoration might vary in their efficacy, and a next step in understanding the efficacy of a Field of Dreams approach is to comprehensively evaluate the interaction between all three scales of partial restoration: spatial, temporal, and ecological.

Resolving the effect of these different scales on restoration success is particularly relevant to systems with the potential for alternative stable states and threshold dynamics. If an ecological system exhibits multiple stable states for a single set of environmental conditions, disturbance can lead to a shift in the system to an undesirable state or ecosystem function with impeded recovery (Beisner et al., 2003). In this case, the unstable threshold represents a target restoration must cross for recovery to occur (Suding and Hobbs, 2009). In the context of the Field of Dreams hypothesis, such a threshold can provide specific partial restoration goals that have to be fulfilled before natural recovery is possible. While the potential for alternative stable states has been identified across terrestrial (James et al., 2013; Ratajczak et al., 2014) and marine (Mumby et al., 2013; Connell et al., 2013; Selkoe et al., 2015) systems (further reviewed in Folke et al. (2004)), establishing whether such states represent prohibited versus slowed recovery is difficult to resolve empirically given challenges over resolving community outcomes at large temporal and spatial scales (Petraitis and Dudgeon, 2004).

A system that exemplifies these multi-faceted components of partial restoration is temperate rocky reefs. Temperate rocky reefs have experienced kelp declines and associated increases in kelp-grazing urchins in several parts of the world (Krumhansl et al., 2016), including southern Australia (Layton et al., 2020) and northern California (Rogers-Bennett and Catton, 2019), motivating novel restoration initiatives (Morris et al., 2020). In addition, temperate rocky reefs can exsit in kelp forest or urchin barren states, which might represent alterantive stable states depending on an array of nonlinear feedbacks (Ling et al., 2015). For example, urchins typically subsist off of kelp blades that detach from extant kelp and drift into the seafloor (“drift kelp”), such that grazing does not cause kelp mortality, especially when predator presence induces cryptic urchin behavior (Harrold and Reed, 1985). However, at low kelp densities, which might arise from environmental disturbances such as heat stress, low nutrients, or storms (Bell et al., 2015), urchin starvation and low predator density can lead to more active kelp grazing, which further increases kelp mortality (Harrold and Reed, 1985). In this sense, outcomes in kelp forests likely arise from a mix of bottom-up and top-down processes Graham et al. (1997); Karatayev et al. (2021); McPherson et al. (2021). Urchins in urchin-dominated barrens can go dormant for prolonged periods of time and at high densities that limit the capacity for kelp to settle. This has lead restoration efforts to focus on urchin removal (Leinaas and Christie, 1996; Watanuki et al., 2010). However, the spatial and temporal extent of urchin removal necessary for kelp recovery is uncertain, and several strategies that extend the ecological scale of restoration, such as kelp reseeding (introducing kelp seeds or juvenile stipes) and outplanting (planting mature kelp stipes), are under exploration (Eger et al., 2020; Morris et al., 2020).

For example, the Sonoma and Mendocino County coastlines of northern California experienced a 95% decline in bull kelp (*Nereocystis luetkeana*) forest coverage (McPherson et al., 2021). These declines occurred due to multiple factors including anomalously warm seawater temperatures between 2014-2016 and nutrient-poor water, that stress kelp and increase purple urchin (*Strongylocentrotus purpuratus*) recruitment (Rogers-Bennett and Catton, 2019; McPherson et al., 2021), and the local extinction of the sunflower sea star (*Pycnopodia helianthoides*), the main natural predator of urchins in this region, due to the sea star wasting disease outbreak in 2013 (Harvell et al., 2019). This decline in kelp coverage has led to the starvation of other herbivores, which has resulted in the closure of the recreational red abalone (*Haliotis rufescens*) fishery and the decline of the commercial red sea urchin (*Mesocentrotus franciscanus*) fishery (Rogers-Bennett and Catton, 2019). This economic impact has accentuated the demand to restore the kelp forest ecosystems in this region. Proposed restoration strategies include urchin removal, kelp reseeding, and outplanting (Hohman et al., 2019). The novelty of these restoration efforts provides high uncertainty on what impact might they have and how they will influence the spatiotemporal patterns of purple urchin density.

In this paper, we use a dynamical population model to explore how the spatial, temporal, and ecological scales of restoration extent influence restoration efficacy in the context of bull kelp restoration in the northern California temperate rocky reefs. To do this, we analyze two metrics for restoration efficacy: the threshold urchin density for natural kelp recovery and the rate of kelp recovery. We evaluate the spatial scale by exploring how varying the portion of the intervened coastline by restoration influences these metrics. We explore the temporal scale by applying the intervention either just at the beginning of the restoration project or through continuous efforts. Finally, we explore the ecological scale by analyzing how reintroducing kelp or reducing urchin density, separately and in combination, influences these metrics.

## 2 Methods

### 2.1 Model overview

In this subsection we present an overview of the model we use to describe the spread dynamics of kelp, and in the following subsection we provide a mathematical formulation of the model. This model follows the distribution of kelp and urchin populations through survival, reproduction, and dispersal over a one-dimensional coastline (Figure 1).

**Figure 1:**
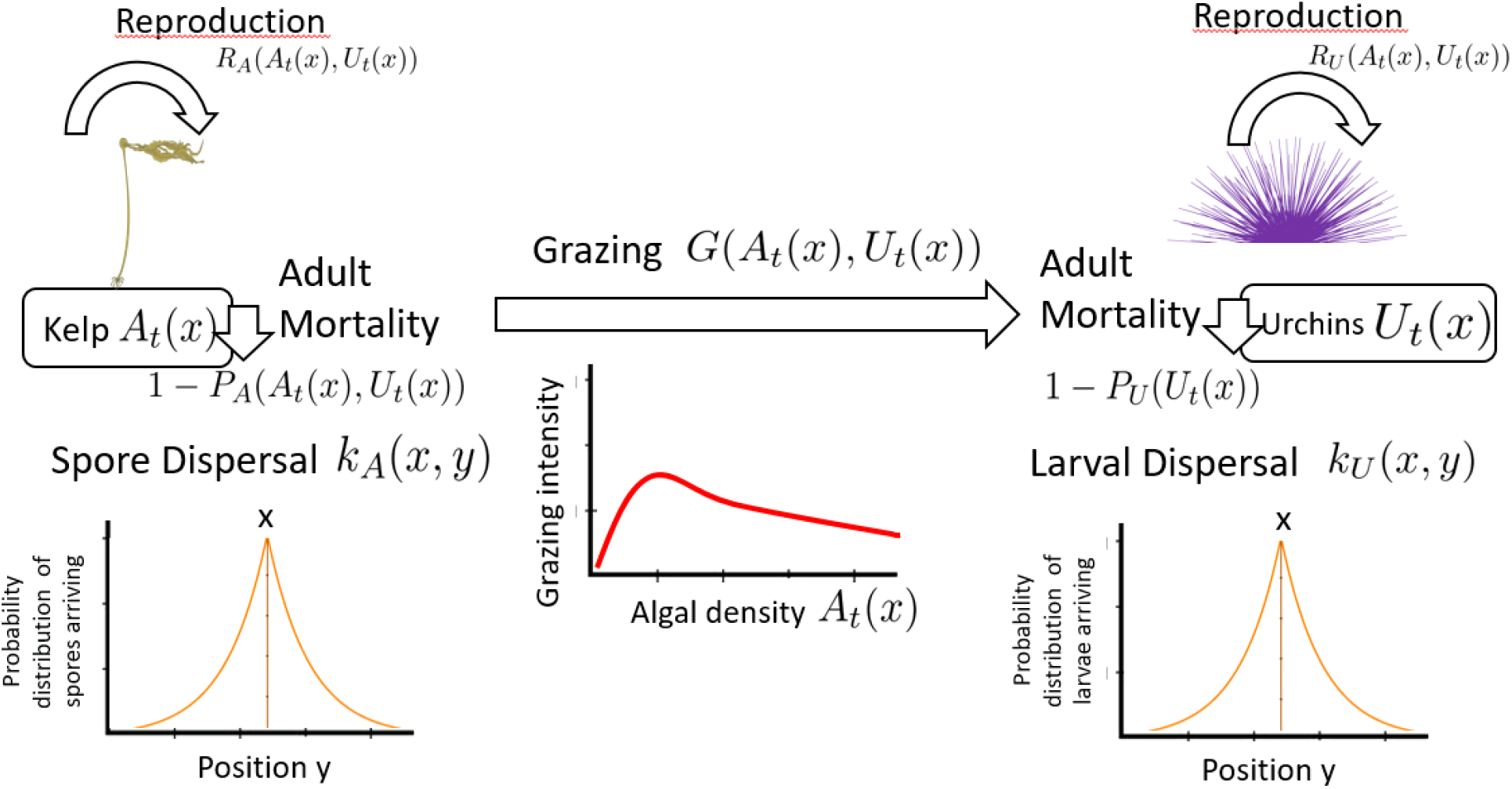
Overview of the dynamics of the model with the respective functional forms of the spore dispersal and grazing intensity. At each time step, a proportion of adults of each species dies off and recruits are produced and dispersed. The grazing interaction affects the spore production of urchins and kelp mortality. Diagram images thanks to Janes Thomas, IAN Image Library (https://ian.umces.edu/imagelibrary/)

At each time step the adults survive with a given probability. We assume that urchin survival is density independent. Kelp survival depends on the grazing intensity by urchins, which depends on both urchin and kelp density. Direct grazing intensity is unimodal with kelp, at first increasing with resource availability and then decreasing at high kelp densities, as might occur due to a switch from active grazing to passive subsistence off of drift kelp (Harrold and Reed, 1985). Adult kelp produces spores at a constant per capita amount, whereas urchins larvae production depends on kelp grazing and drift kelp consumption. Spores and larvae then disperse through the coastline and a fraction of them settle and become adults. In line with observations of urchin adult movement on the scale of a few meters (Dumont et al., 2006), kelp seeds and zoospore movement on the scale of tens of meters (Dobkowski et al., 2019), and urchin larval movement on the scale of kilometers (Largier, 2003), we assume that adult urchin movement is significantly smaller than dispersal of the kelp and urchin juvenile stages, and thus neglect any adult urchin movement.

We vary the amount of urchin removal, kelp reseeding, and kelp reintroduction over a range of spatial and temporal extents. We focus on these interventions, and do not include predator reintroduction as well, for two reasons. First, research into the feasibility of seastar reintroduction as a restoration intervention for our focal system of the California north coast is still in development and at the stage of lab tests (J. Hodin, personal communication), while urchin removal is underway (Ward et al., 2022) and kelp reseeding and reintroduction are undergoing field tests (B. Hughes, personal communication). Second, an ongoing question for the role and timing of predator reintroduction is whether and, if so, how much predators will seek out and gain energy from non-barren urchins (i.e. urchins in recovered kelp stands with enough kelp consumption to support gonad production), such that predator reintroduction might be more effective if it occurs after interventions that increase kelp density. Given these considerations, current restoration efforts, which we seek to inform, are focused on urchin removal, kelp reseeding, and kelp introduction (Hohman et al., 2019).

### 2.2 Model

Our model combines the ecological dynamics of Karatayev et al. (2021) with the spatial dynamics of Kanary et al. (2014). We consider populations of kelp and urchins cohabit in a one-dimensional coastline Ω. Our model follows kelp (*A_t_*(*x*)) and urchins (*U_t_*(*x*)) through time *t* and space *x*. At each time step and for each species *i* (*i* = *A* for kelp and *i* = *U* for urchins), the adults survive to the next step following the function *P_i_*(*A_t_*(*x*), *U_t_*(*x*)) and adults produce recruits according to a function *R_i_*(*A_t_*(*x*), *U_t_*(*x*)). The recruit survive to the next step following a function *S_i_*(*A_t_*(*x*), *U_t_*(*x*)) and disperse from their source following a kernel *k_i_* in an integrodifference equation framework. Combining these dynamics, the populations for algae and kelp at the next time step follow:

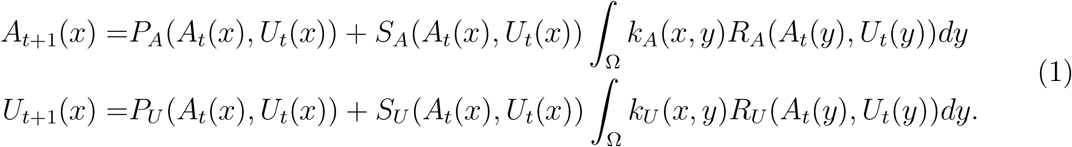

Adult kelp has a natural survival probability in absence of urchin grazing given by *δ_A_*, which implies a mean lifespan of 1/*δ_A_*. In addition, kelp survival depends on urchin grazing, which we model as a Holling’s “Type IV” functional response *G*(*A_t_*(*x*), *U_t_*(*x*)). This functional response phenomenologically represents a behavioral shift from active to passive grazing with increasing kelp and can lead to two alternate stable states: a kelp-dominated state (kelp forest) and an urchin-dominated state (urchin barren) (Karatayev et al., 2021), which occurs under our parameterization (Appendix S1). Urchins graze kelp holdfast with a base attack intensity *γ_A_*. Given a maximum grazing consumption at 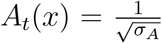, adult kelp survival is:

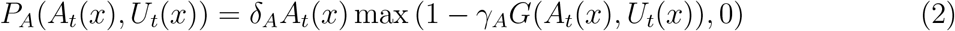

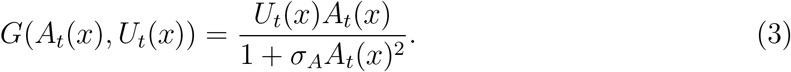

We assume that kelp produces a constant per capita number of spores *R*, which gives *R_A_*(*A_t_*(*y*), *U_t_*(*y*)) = *RA_t_*(*y*). Kelp spore survival and recruitment depends on two factors: the probability of spores settlement and urchin predation. Settlement is density-dependent with a saturating, Beverton-Holt-type function given the maximum kelp population at a given location *x* of 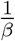. In addition, we assume that urchins graze recently settled kelp stipes before they can grow to a mature sporophyte with a per capita probability *γ_S_*. Then, the survival of kelp spores is

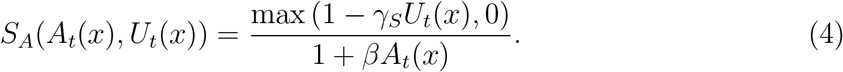

Urchin survival occurs with a constant probability *δ_U_*, which gives us *P_U_* (*A_t_*(*x*), *U_t_*(*x*)) = *δ_U_ U_t_*(*x*). Urchin larval production arises from two sources. First, urchins gain energy through direct grazing, proportional to Equation 3 with a proportion constant *γ_U_* (Neubert et al., 1995). Second, urchins gain energy for larval production through drift kelp consumption at a constant proportion *ε* of the kelp available at each location *x*. Both *γ_U_* and *ϵ* encapuslate conversion of energy gained from kelp consumption into larval production and survival such that *S_U_* (*A_t_*(*x*), *U_t_*(*x*)) = 1. Combining both sources of energetic gain, the total urchin larval production is

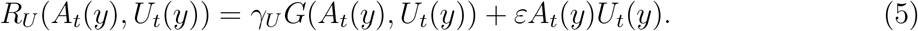

Finally, we model both dispersal kernels as Laplace kernels with mean dispersal distance for each species *i* 1/*a_i_* given by the equation (Lockwood et al., 2002):

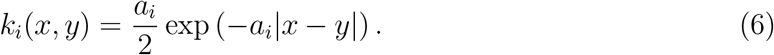

Note that, with constant and homogeneous kelp natural mortality *δ_A_*, kelp spore production *R*, and urchin production *γ_U_* and *ε*, we focus on kelp-urchin interactions and ignore the role of seasonal and variable environmental conditions in driving kelp and urchin dynamics. We make this simplifying assumption because of our focus on restoration decisions concerning the choice of urchin removal and kelp reintroduction interventions at different spatial extents and temporal scales. Informing the additional (and important) restoration decisions of optimal location and timing of restoration interventions, not under consideration here, would require model extensions that account for spatially heterogeneous and temporally stochastic environmental drivers such as nutrients, light, and wave disturbance that can influence kelp dynamics (Graham et al., 1997; Karatayev et al., 2021), as well as the stochasticity and seasonality of urchin reproduction (Okamoto et al., 2020; Cochran and Engelmann, 1975).

### 2.3 Parameter estimation

We fit the model without interventions to kelp and urchin distribution data in the Sonoma-Mendocino coast. We compile yearly kelp coverage data from the dataset of McPherson et al. (2021) with the yearly urchin data of Reef Check (ReefCheck, 2020). We estimate all parameters except *β* using the 2007 and 2008 data (Figure 2). We identify two regions with available urchin data, which correspond to the coasts of Little River and Timber Cove.

**Figure 2:**
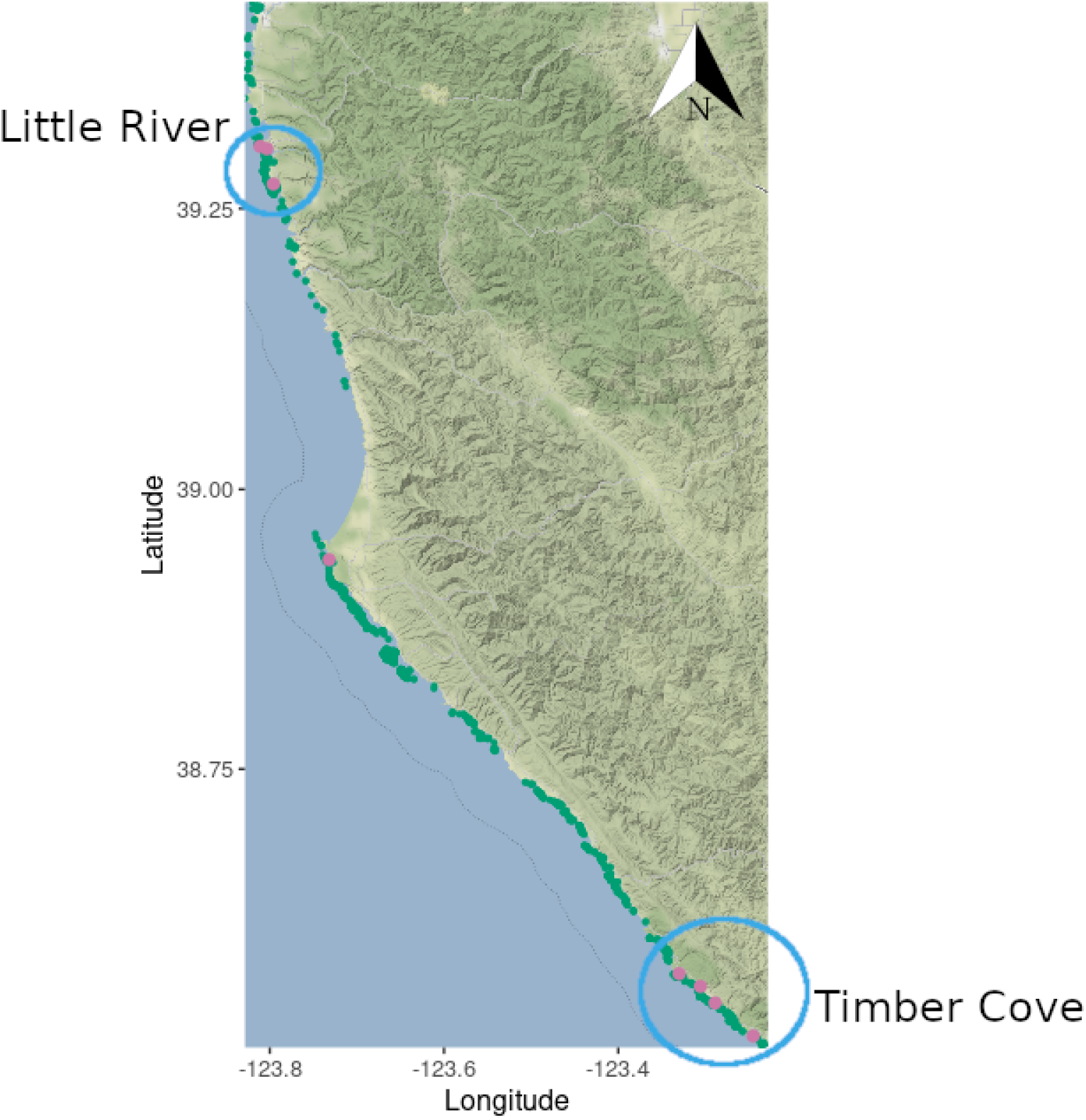
Kelp (green dots) and urchin (purple dots) data available for 2007-2008 in the Sonoma and Mendocino counties of California. Notice the highlighted regions, circled in blue, where urchin data is available. These regions correspond to Little River (top circle) and Timber Cove (bottom circle). Notice the purple dot in the middle of the map corresponds to a single spatial point, which makes our spatial analysis unfeasible.

We estimate the parameters using the Approximate Bayesian Computation (ABC) method, implemented using the EasyABC package in R (Jabot et al., 2013). In order to reduce estimation errors due to possible parameters correlations, we will implement the ABC algorithm with Metropolis-Hastings sampling, implemented in the EasyABC package as the Marjoram method and described in Wegmann et al. (2009).

Using the ABC algorithm, we start our simulations taking the initial conditions *A*_0_(*x*), *U*_0_(*x*) being the distribution for each of the regions at the 2007 measurement. We initialize our ABC algorithm with uniform prior distributions for each of the parameters in the range presented in Table 1. We then run the model for a sampled combination of parameters for a year (where each time step *t* corresponds to 1 month) and compare the obtained kelp distribution *A*_12_(*x*) with the distribution at the 2008 measurement. We compare these distributions through using RMSE as our summary function. In other words, if *A*_12_(*x*) is the 2008 distribution in the given region, we find combinations of parameters that minimize:

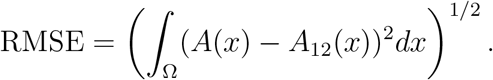

**Table 1:** Description of each of the parameters of the model.

| Parameter | Description | Range of Possible Values Explored | Best-fit Value |
| --- | --- | --- | --- |
| $\delta_A$ | Survival probability of adult kelp | [0, 1] | 0.510 |
| $\gamma_A$ | Grazing intensity of urchins on kelp | [0, 1] | 0.101<br>kelp m<sup>-2</sup><br>urchins<sup>-1</sup> |
| $\sigma_A$ | Inversely determines kelp density at maximum urchin grazing | [0, 100] | 15.475<br>kelp m<sup>-2</sup> |
| $\gamma_S$ | Probability of juvenile kelp stipes being grazed by urchin | [0, 100] | 0.743<br>urchins<sup>-1</sup> |
| $\beta$ | Inverse of maximum kelp density | Estimated without ABC (see text) | 2.42 kelp m<sup>-2</sup> |
| $a_A$ | Inverse of mean dispersal distance of kelp | [0, 100] | 16.137<br>m<sup>-1</sup> |
| $R$ | Per capita spores production of kelp | [0, 10] | 5.500 |
| $\delta_U$ | Survival probability of urchins | [0, 1] | 0.312 |
| $\gamma_G$ | Urchin production from direct kelp consumption | [0, 10] | 4.956<br>urchins |
| $a_U$ | Inverse of mean dispersal distance of urchin | [0, 100] | 93.586<br>m<sup>-1</sup> |
| $\varepsilon$ | Urchin production from kelp consumed by urchins as drift kelp | [0, 10] | 9.484<br>kelp m<sup>-2</sup> |
| $\mu_U$ | Intensity of urchin removal relative to natural urchin mortality | [0, 1] | |
| $\mu_S$ | Intensity of kelp seeding relative to per capita spores production | [0, 1] | |
| $\mu_A$ | Intensity of kelp outplanting relative to natural kelp mortality | [0, 1] | |
| $\eta$ | Length of region to apply restoration efforts around a kelp oasis | [0, 100] | |
| $A_c$ | Critical kelp density to identify where to apply restoration efforts | [0, 1] | |
| $A_0$ | Initial kelp density at the kelp oasis | [0, 5] | |
| $U_0$ | Initial urchin density at the coastline | [0, 100] | |

While this approach ignores the seasonal nature of kelp and urchin recruitment as well as kelp mortality from winter storms (Ebert et al., 1994; Springer et al., 2010), in the absence of monthly data that would allow model-fitting to seasonal processes, it does capture the year-to-year dynamics that match the time scale of the data.

To estimate the *β* parameter that inversely determines the kelp recruitment saturation level, we perform a linear regression at each point in space in the kelp distributions from 2004 to 2009 and fit it to a Beverton-Holt model (Beverton and Holt, 1993). This procedure allows us to make use of the higher availability of kelp data, and reduce the number of parameters our ABC procedure has to estimate. We then use the distribution of maximum densities as our distribution for 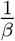.

### 2.4 Control strategies and model analysis

To explore what control strategies promote the spread of kelp, we follow the restoration focus of the Sonoma-Mendocino Bull Kelp Recovery Plan (Hohman et al., 2019). This Recovery Plan focuses in implementing several restoration strategies near kelp “oases”, i.e. patches of extant kelp, to try to enhance kelp expansion to nearby regions. We explore three restoration strategies around these oases: urchin removal, kelp re-seeding, and kelp outplanting.

We initialize our simulation with initial condition of kelp *A*_0_(*x*) be 0 everywhere except at a starting oasis of length *L*, in which we start with kelp at an initial kelp density *A*_0_; the urchins initial density is *U*_0_ throughout the coastlines Ω. To identify target restoration locations as kelp density (and therefore the location of oases) changes through time, at each time step *t*, we define the region where restoration efforts are applied as the set of all locations centered around *x* with length *η* where the kelp density surpasses a critical density *A_c_*. This gives us a function *δ_t_*(*x*) to indicated the presence or absence of restoration actions given by

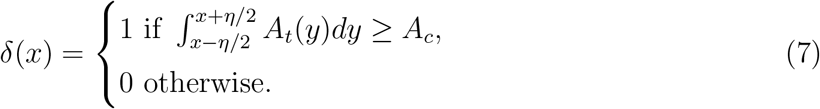

Multiplying implementation presence *δ_t_*(*x*) by the control intensity of urchin removal *μ_U_*, kelp seeding *μ_S_*, or kelp outplanting *μ_A_*, provides the terms for modifying the survival probability of adult urchins, spore production of kelp, and survival probability, respectively, when implementing restoration, yielding

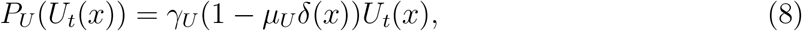

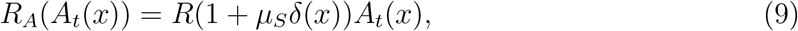

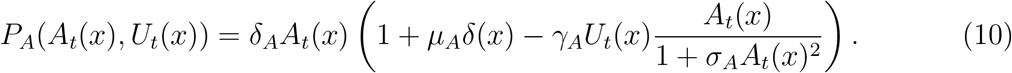

We explore the temporal scale of restoration through two scenarios: first we implement a short time scale restoration effort by varying the initial densities *A*_0_ and *U*_0_ and setting long-term control *μ_i_* = 0 for function *i* = *U, S, A*. This corresponds to the case in which partial restoration efforts are performed at the start of the project, and then natural processes (e.g. succession) might eventually achieve restoration goals. Second, we implement a long time scale restoration effort where the initial kelp and urchin densities *A*_0_ and *U*_0_ are fixed, and we vary the restoration effort intensity *μ_i_*. For these simulations, we set the initial kelp density *A*_0_ = 1/*β* within the oasis (and zero elsewhere) and initial urchin density *U*_0_ to 95% of the threshold value that the system must cross for kelp recovery. Then, for ongoing restoration intervention, we explore a range of intensities in terms of a percentage increase in urchin mortality (urchin removal), kelp spores (seed outplanting), and adult kelp (kelp outplanting). In each case we explore weak (10%), moderate (40%), and strong (70%) intensities.

In both temporal scale scenarios, we explore the effect of different spatial scales. For the short time scale scenario, we explore the spatial scale of restoration by varying the length of the initial oasis *L*. In the long time scale scenario, we explore the spatial scale of restoration by varying the size of the region with a control effort *η*. Finally, we explore the ecological scale of restoration by comparing scenarios with only a single control strategy or a combination of strategies (urchin removal, kelp re-seeding, or kelp outplanting) at varying intensities.

We evaluate these scenarios using two metrics. Our first metric is the maximum initial urchin density at which kelp can spread (hereafter the urchin threshold), which represents the restoration effort necessary for eventual recovery to take place. The second metric is the kelp recovery rate (hereafter spread rate). To calculate the spread rate, we run the system for 12 time steps (months) and, at each time step, calculate spread extent as the distance from the starting point *x* = 0 to the point *x* where there is a significant amount of kelp coverage is more than 1% of kelp coverage. We then calculate the spread rate as the slope of the linear regression of spread extent versus time. We choose months as our time scale to explore the dynamics of our system through the span of a single year, which allows us to see the short-term effect of the different restoration strategies, while also accounting for the annual nature of bull kelp, where factors not modelled, such as storm disturbance, might further affect kelp survival at the end of our time horizon.

In order to quantify the relative effect of different processes and management levers on the urchin threshold and spread rate, we perform a global sensitivity analysis of all parameters in the model, based on the procedure by Harper et al. (2011). We first sample 2000 combinations of parameters from the posterior distributions obtained from the parameter estimations and calculate the urchin thresholds and spread rates for each combination. We then construct a random forest using the R package randomForest (Liaw and Wiener, 2002), with the parameters of our model as predictors and the urchin threshold or spread rate as the target function. The randomForest package provides an importance metric for each predictor, which indicates how frequently that predictor served a breakpoint in the random trees of the forest.

## 3 Results

In our model parameterization, the ABC presented high uncertainty in all of the estimated parameters; we focus on our global sensitivity analysis influence of the different parameters on the model outcomes. We only show the results for the parameter estimates of the Timber Cove region, using the best-fit values presented in Table 1. We do this because the ABC of Timber Cove provided better posterior distributions than that of Little River, and thus less uncertainty. See Appendix S2 for the posterior distributions for both regions.

### 3.1 Urchin threshold

Given a short-term, one-time urchin removal, the threshold urchin density necessary for kelp recovery increases if, in addition to urchin removal, further kelp is planted (Figure 3). This occurs because increasing kelp density lowers the grazing intensity of urchins due to the behavioral feedback in the Type IV functional response of urchin grazing. Accordingly, an increase in *σ_A_* (the parameter which determines the strength of the behavioral feedback) increases the threshold urchin density for kelp recovery. In the context of the Field of Dreams hypothesis, kelp natural recovery can be feasible after removing urchins below a certain threshold. Increasing the ecological scale of restoration through including kelp outplanting (i.e., increasing initial kelp density) increases this threshold, which reduces the intensity of urchin removal efforts required to ensure kelp recovery.

**Figure 3:**
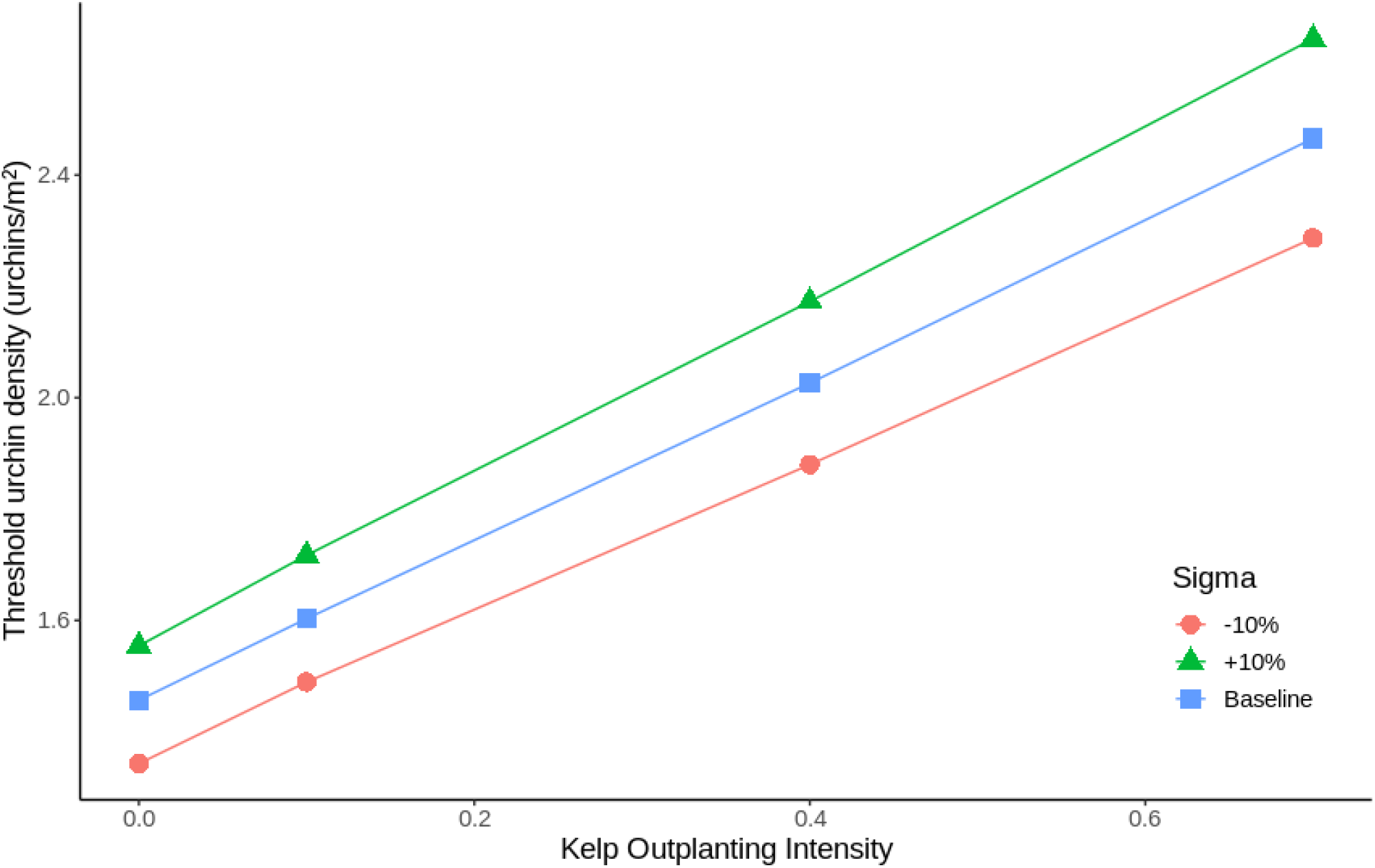
The threshold urchin density necessary for kelp recovery as a function of kelp outplanting intensity. Different lines represent different values of *σ_A_* (inversely determines the kelp density of maximum urchin grazing in the Type IV functional response) changed by 10% from its baseline value.

Our global sensitivity analysis (Figure 4) confirms the main factor that affects this threshold urchin density for kelp recovery is the urchin grazing activity (described by the conversion of kelp to urchins *γ_A_* and the kelp grazing inhibition parameter *σ_A_*). Specifically, the threshold urchin density is greater for slower urchin grazing (lower *γ_A_*) and a lower peak value for direct kelp grazing (lower *σ_A_*). In addition, the threshold urchin density is higher for a higher urchin natural mortality rate (lower urchin survival probability *δ_U_*), higher initial kelp density (higher initial kelp density *A*_0_), and with a lower kelp natural mortality (higher kelp survival probability *δ_A_*).

**Figure 4:**
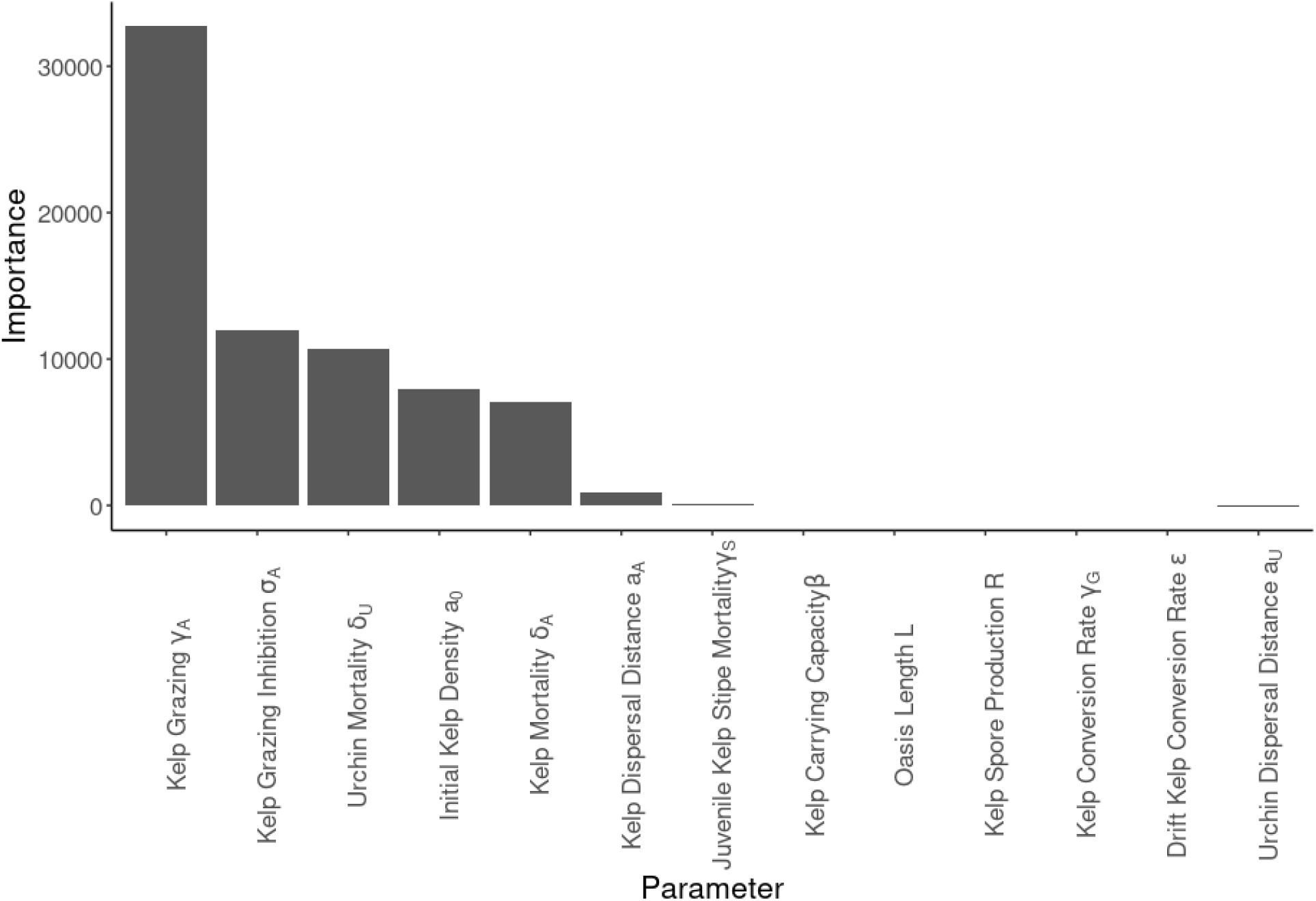
Importance ranking of the parameters of the Model 1 from the global sensitivity analysis of the threshold urchin density necessary for kelp recovery. See Table 1 for more detailed parameter definitions.

### 3.2 Kelp spread rate

Given an initial urchin removal below the threshold value required for recovery, kelp spread rate increases with expanding interventions across ecological scales more than expanding over spatial or temporal scales. Under our baseline parameterization, ongoing kelp seeding enhances kelp recovery rate, while long-term, ongoing kelp outplanting or urchin removal do not (Figure 5). However, the relative effect of other strategies is sensitive to initial kelp density. With double the initial kelp density, further kelp outplanting increases kelp recovery rate (Figure 6a), while with triple the initial kelp density, ongoing urchin removal has a bigger effect on kelp recovery rate (Figure 6b). The greater sensitivity to initial kelp density for ongoing urchin removal and kelp outplanting, as compared to kelp reseeding, is likely due to the nonlinear (Type IV) feedback between urchin grazing and extant kelp as compared to the linear (Type I) feedback between urchin grazing and kelp seeds. These different dynamics lead to a different influence of the control strategies that directly affect the local kelp-urchin interaction.

**Figure 5:**
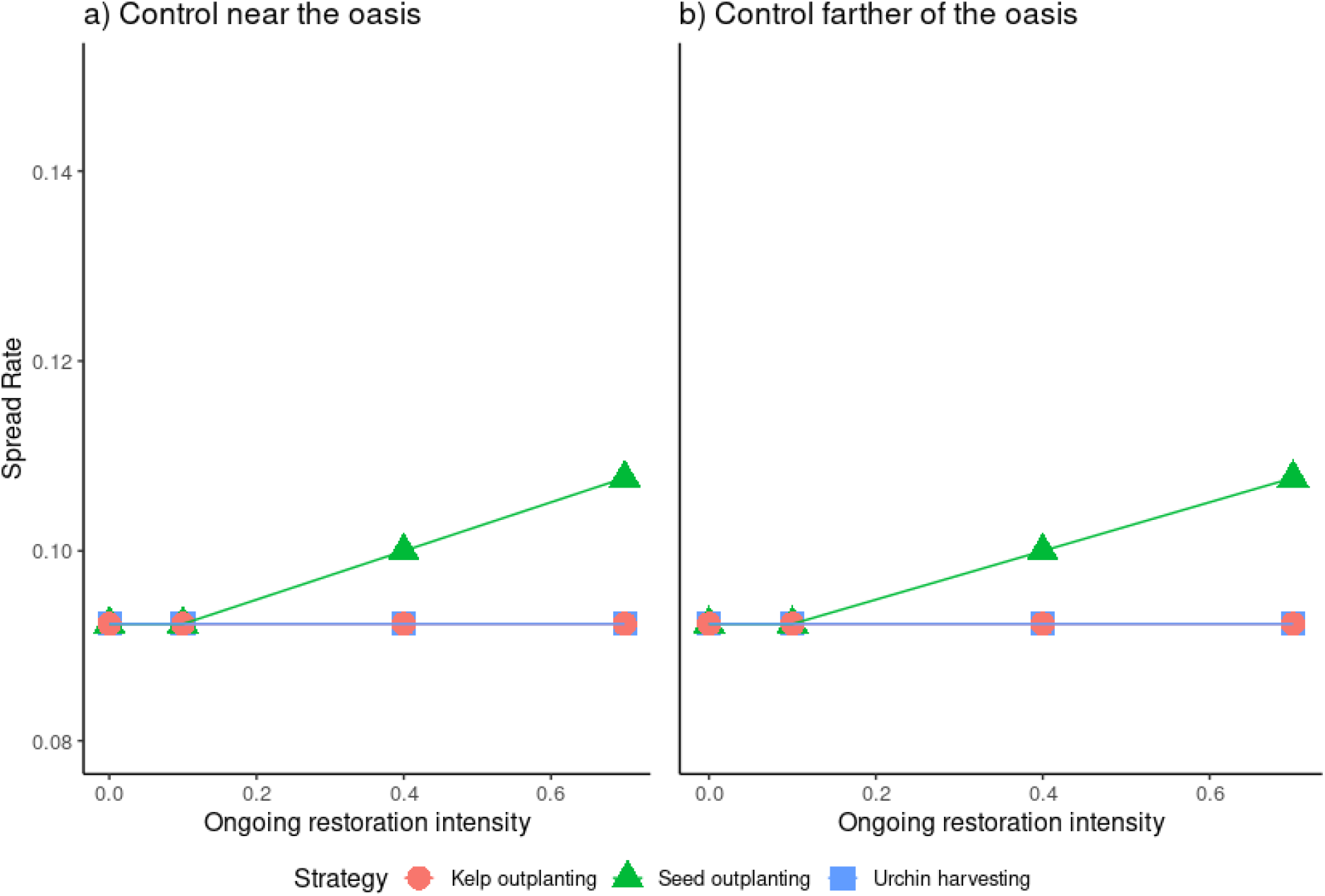
Kelp spread rate under different restoration strategies with increasing intensity following urchin removal below the threshold value necessary for kelp recovery and with an initial kelp density 1/*β*. Each line represents a different strategy: kelp outplanting in red circles, kelp seeding in green triangles, and sustained urchin harvest in blue squares. Panel a) shows ongoing restoration efforts near the kelp oasis (*η* = 1), and panel b) shows ongoing restoration efforts across a wider region of the coastline (*η* = 10).

**Figure 6:**
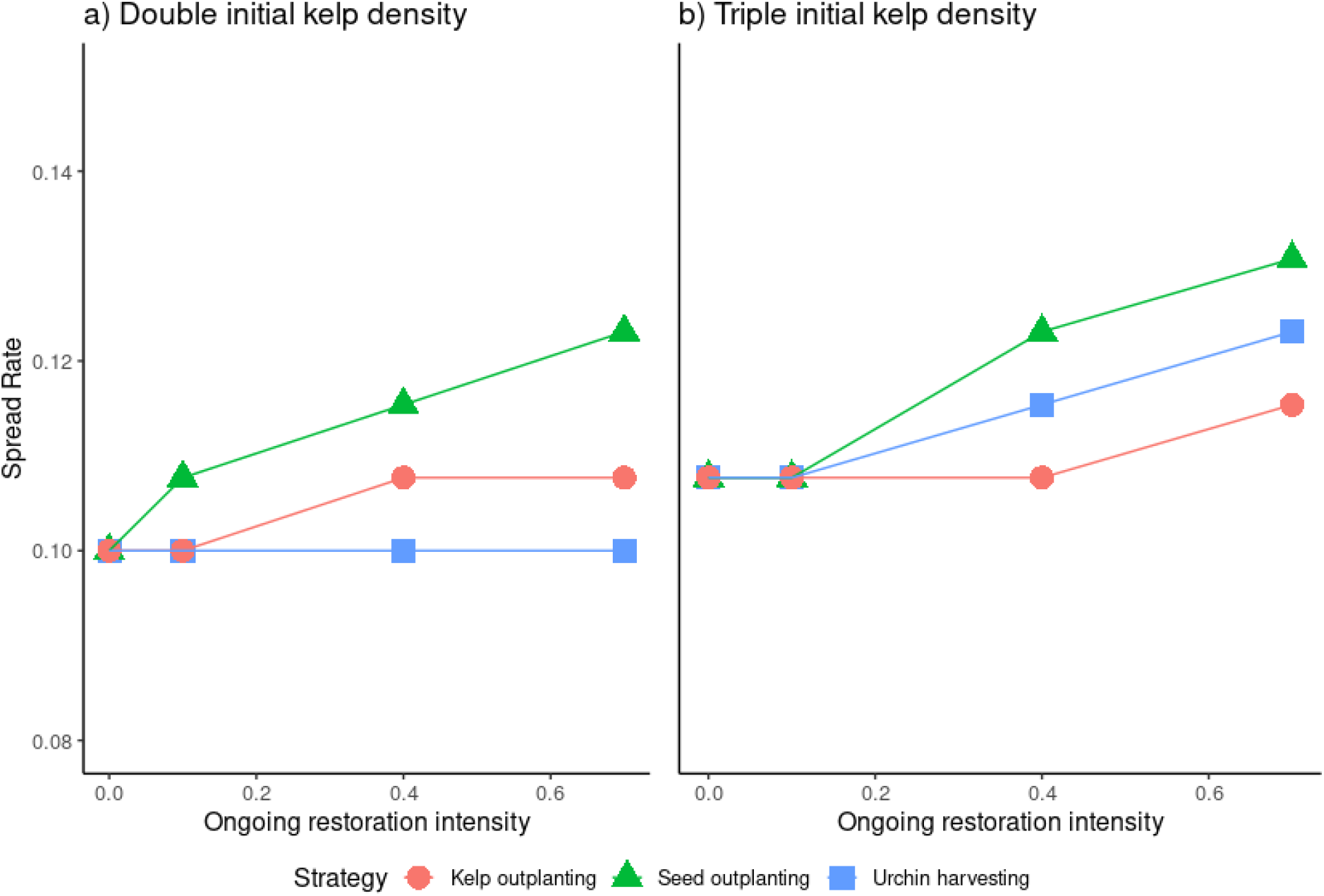
Kelp spread rate under different restoration strategies with increasing intensity following urchin removal below the threshold value necessary for kelp recovery with a varying initial kelp density. Each line represents a different strategy: kelp outplanting in red circles, kelp seeding in green triangles, and sustained urchin harvest in blue squares. Panel a) shows ongoing restoration efforts with double the initial kelp density in the oasis compared to the default of 1/*β*, and panel b) shows ongoing restoration efforts with triple the initial kelp density.

For the spatial scale of restoration, increasing the extent of ongoing restoration efforts does not affect the rate of recovery of kelp (compare panels a) and b) of Figure 5). This suggests that kelp recovery is mainly determined by the local urchin grazing intensity, and extending restoration efforts to regions of the coastline with reduced kelp densities, where urchin grazing is stronger, will not affect kelp recovery rate. For the temporal scale of restoration, increasing the initial kelp density and decreasing the initial urchin density through a more intense partial restoration effort at the beginning enhance kelp recovery more than ongoing restoration efforts (compare the spread rates in Figure 7 to those in Figures 5 and 6). Increasing kelp density and reducing urchin density near the kelp oasis provides better conditions for kelp survival when interacting with the urchins, which further enhances kelp recovery rate. Overall, the combination of increasing initial kelp density, reducing initial urchin density, and implementing an ongoing kelp seeding leads to the fastest kelp recovery.

**Figure 7:**
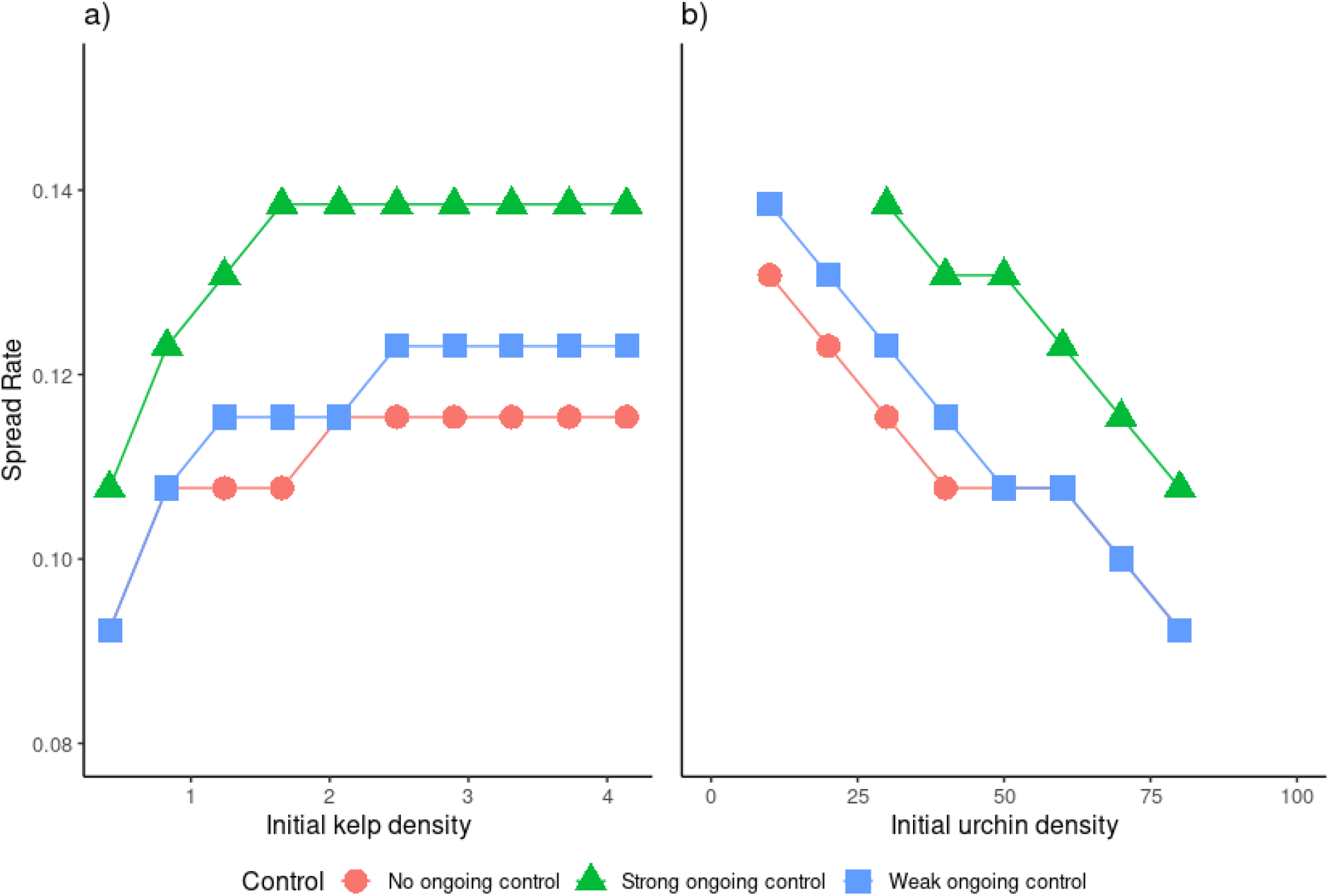
Kelp spread rate under different initial conditions for partial restoration efforts at the initial (short-term) restoration stage. Each line represents a different ongoing restoration effort in terms of kelp seeding: no ongoing restoration effort in red circles, strong ongoing seeding in green triangles (*μ_S_* = 0.7), and weak ongoing seeding in blue squares (*μ_S_* = 0.1). Panel a) shows the change in spread rate as the initial kelp density varies, and panel b) shows the change in spread rate as the initial urchin density varies.

The primary role of short-term restoration efforts is further evident in the global sensitivity analysis of the spread rate (Figure 8), where initial kelp and urchin densities (*A*_0_ and *U*_0_ respectively) have a higher impact over the spread rate than ongoing restoration efforts (*μ_i_* for *i* = *U, A, S*). Therefore, both natural local conditions that lead to higher kelp coverage and lower urchin densities after a marine heatwave, as well as interventions to increase kelp density and decrease urchin density, have a strong impact on overall spread rate. When comparing the importance of the parameters for the threshold urchin density (Figure 4) and kelp recovery rate (Figure 8), we observe that parameters such as size of the oasis (*L*) and mean dispersal distance of kelp (*a_A_*) play a role on kelp recovery once urchin density is below the threshold necessary for recovery. Intuitively, a higher mean dispersal distance of kelp seeds (*a_A_*) leads to a faster spread (Figures 3–5 in Appendix S3), especially for the strategy of seed outplanting. However, note that the relative efficacy of the different restoration strategies remain unchanged for different values of mean dispersal distances of kelp seeds. In addition, the lower importance value of kelp dispersal distance compared to parameters related to urchin grazing indicates that local kelp-urchin interactions have a greater influence on kelp spread rate than kelp dispersal. While increasing the size of the initial kelp oasis (*L*) through kelp outplanting enhances kelp recovery rate, the spatial scale of ongoing restoration efforts (*η*) has a minimal impact over kelp recovery rate.

**Figure 8:**
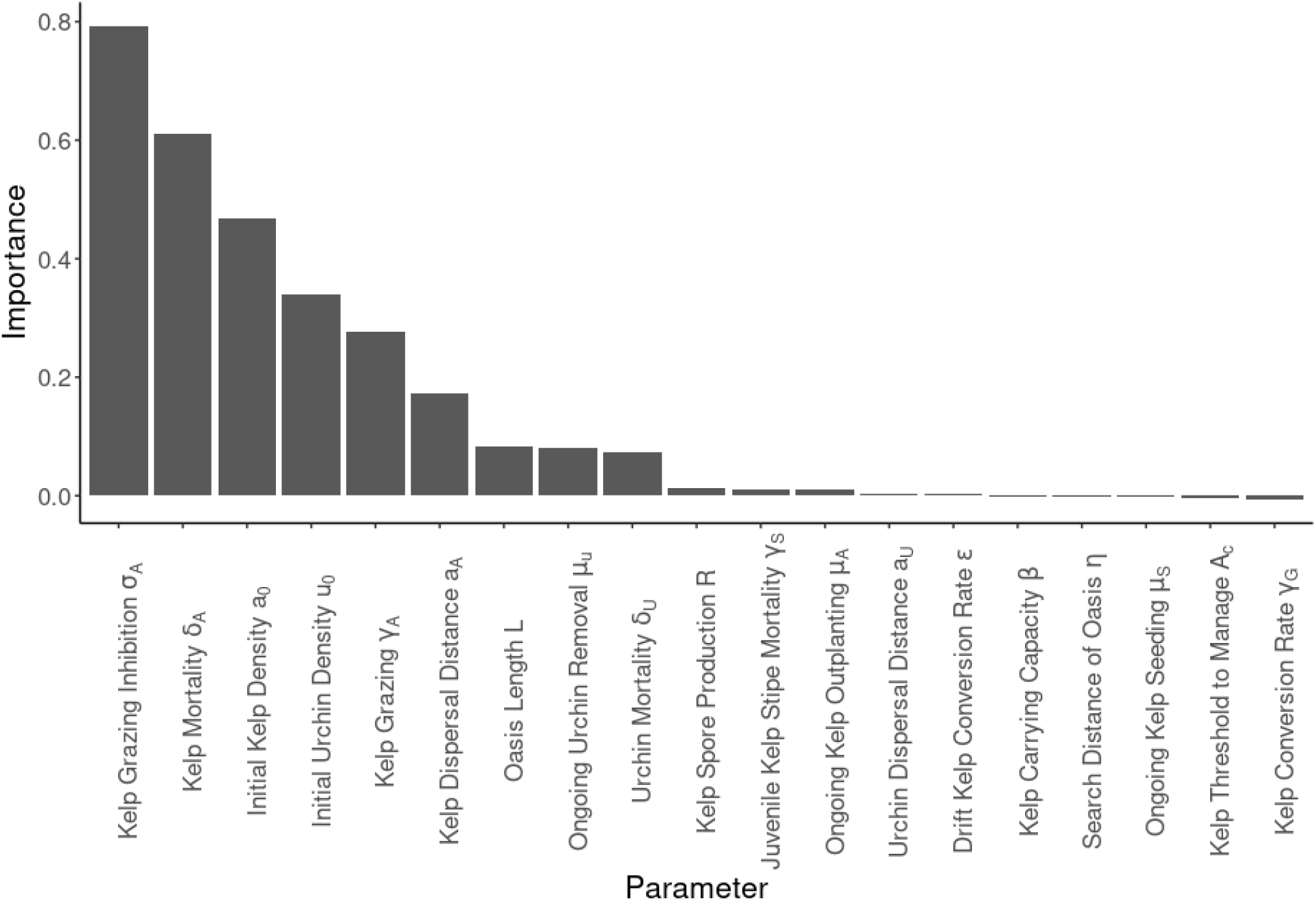
Importance ranking of the parameters of the Model 1 from the global sensitivity analysis of the kelp recovery rate. See Table 1 for more detailed parameter definitions.

## 4 Discussion

In our model of kelp restoration, scaling up ecologically on restoration efforts can have a bigger effect in restoration success than scaling up spatially or temporally. One of the key factors in determining if kelp recovery will be possible is the threshold urchin density, which our model suggests is mostly determined by local interactions. Because we incorporated kelp-urchin grazing feedbacks that can drive alternative stable states, kelp recovery does not occur in our model unless urchin density is below a certain value (Figure 3). This threshold increases as kelp density increases, i.e. kelp outplanting reduces the amount of urchin removal necessary for recovery. In addition, improving the initial conditions through an increase in kelp density or decrease in urchin density at an early stage can enhance kelp recovery rate more than ongoing restoration efforts (compare Figures 5 and 7). This suggests that a Field of Dreams approach can apply in kelp forest restoration, as in our model enhancing the ecological conditions at a short temporal scale and small spatial scale has more impact than distributing restoration efforts through a longer period of time or to a greater spatial extent.

The theoretical results of the different restoration outcomes are consistent with what has been observed in kelp restoration efforts. In recent restoration efforts in our focal system of Northern California, sites where urchin removal has been implemented below a threshold density of 2 urchins/m^2^ had seen significantly higher kelp density compared to sites without urchin removal (Ward et al., 2022). While these restoration sites and regions of improved recovery cover a much smaller spatial scale compared to original loss (compare Rogers-Bennett and Catton (2019) to Ward et al. (2022)) due to resource limitation, they can facilitate kelp recovery in targeted locations of economic importance (e.g. near ports) to coastal stakeholders such as the fishing and diving communities. In Southern California, sea urchin removal can increase the success of kelp reseeding Ford and Meux (2010). Beyond California, Layton et al. (2020) found that urchin removal and kelp outplanting had been successful restoration methods for increasing kelp density in different regions of the Australian coastline. In the case of kelp reintroduction on the coast of Tasmania, Sanderson (2003) found greater success of kelp outplants in areas with urchin removal.

This predominant role of early restoration efforts also parallels empirical findings in systems beyond the kelp system modeled here. For example, in Lee et al. (2006) restoring the habitat of anurans (wetlands) at a small spatial and temporal scale was enough to allow eventual recovery of community composition and diversity of amphibians. In Cahall et al. (2013), thinning of the forest at an early stage led to an increase in the density of certain bird populations compared to unthinned (Figure 7). These cases where a Field of Dreams approach is successful have in common that the habitat quality is one of the main limiting factors of restoration success. In the case of kelp forest restoration, a suitable habitat is determined by the active grazer density (purple urchins in our study system).

The predominance of early restoration efforts, and greater efficacy of ecological scaling up over spatial or temporal scaling up in restoration efforts, arises, in part, because of the threshold dynamics in our model with alternative stable states. As described above, with these threshold dynamics, once initial restoration passes the threshold, which depends on both urchin and kelp densities, then natural recovery can occur. The potential for alternative stable states arises from our Type IV (unimodal) functional response in urchin grazing to kelp density, which might occur due to an urchin grazing behavioral shift from active to passive grazing due to greater subsistence on drift kelp or cryptic behavior with higher densities of kelp and associated urchin predators (Karatayev et al., 2021). Accordingly, our global sensitivity analyses indicates that the parameters that shape urchin grazing response to kelp density comprise the main driving factors of both the possibility of recovery and its rate. If, in reality, this feedback between kelp density and urchin density is not strong enough to drive alternative stable states, we would expect a reduction in the overall role of active grazing behavior and a potential increase in the role of greater spatial and temporal scales of restoration efforts. That said, our best-fit model did find a strong enough feedback between kelp density and urchin grazing for alternative stable states to occur, and McPherson et al. (2021) provide empirical support of a significant role of urchin grazing in kelp decline on the California north coast, where including urchin grazing into a partial least squares regression analysis doubled the variability in the yearly data of kelp coverage explained by the model. In addition, as noted above, data from restoration efforts indicate that urchin removal can increase restoration success in an number of kelp systems spanning California and Australia (Ward et al., 2022; Layton et al., 2020; Sanderson, 2003; Ford and Meux, 2010). Further evidence for a role for urchin densities in kelp dynamics include rapid kelp recovery following urchin mass mortality in southern California (Williams et al., 2021) and rapid kelp declines following urchin range expansions in Tasmania (Ling, 2008). In Tasmania, both theory and data suggest that alternative stable states between urchin barrens and kelp forests affect recovery success, analogous to our model (Marzloff et al., 2016; Ling, 2008; Johnson et al., 2017).

Although increasing the temporal scale of restoration has a smaller effect on the ability of kelp to recover in our determinisitc simulations, an increase in the temporal scale of restoration can buffer against the potential for short-term restoration failure from caused by extreme, stochastic events (Reich and Lake, 2015). In the case of kelp, an increase in the likelihood of marine heatwaves can lead to potential die-offs of kelp and increases in urchin recruitment, which might bring urchin density above the threshold and restrict kelp recovery (Rogers-Bennett and Catton, 2019). This potential for restoration failure due to environmental stochasticity has been noted in amphibian (Dodd and Seigel, 1991) and plant (Dalrymple et al., 2012) reintroductions, and has been observed in coral reef restoration failing due to hurricane activity (Bowden-Kerby, 2001).

Finally, partial restoration efforts leading to longer-term, larger scale recovery will depend on the spatial extent of the dispersal and the temporal scale of generation time of the ecological components. For example, long river systems may require a timeframe in the scale of decades to reestablish their hydrological dynamics (Tockner et al., 1998). In addition, active interventions may reduce the impact of ecological traps produced by restoration efforts (Hale and Swearer, 2017). For example, Severns (2011) found the butterfly *Lycaena xanthoides* to selectively oviposit more frequently and with more eggs in seasonally-flooded habitats with lower egg survival, as compared to adjacent non-flooded habitats where a tall invasive grass obscured native plants. Averting this accidental cue to poor ovipositing habitat would likely require scaling up of restoration to also incorporate invasive species removal in non-flooded habitats. More generally, active management approaches to account for ecological traps may include changing the behavior of the animals by removing cues or habituating the animals to ignore the cues provided by ecological traps Hale and Swearer (2017).

### 4.1 Management implications

In our model, the most effective restoration approach for kelp forest in northern California is a combination of reduction of purple urchin density through urchin removal and increase of bull kelp density through adult kelp outplanting at an early stage of the restoration project. The role of any ongoing restoration efforts, including further urchin removal and kelp outplanting, as well as kelp seeding was highly sensitive to initial kelp density (Figures 5 and 6), which supports targeting such efforts around extant kelp “oases”. In the absence of any oases, kelp recovery might further rely on initial kelp reseeding or outplanting restoration interventions, depending on the potential for a spore bank as discussed in the “Model limitations” section below. Ongoing restoration efforts might play a greater role if initial removals are insufficient to pass the threshold for kelp recovery or, as noted above, future extreme climate events might disrupt restored populations.

Urchin removal is a technique that is already being applied in the northern coast of California (Hohman et al., 2019). Our results suggest that these efforts are more likely to be successful when complemented with kelp outplanting. Transplantation of *N. luetkeana* has been successfully applied further north in the coast of Washington, where transplanting of juveniles of natural populations was more successful than cultured kelp (Carney et al., 2005). The effectiveness of outplanting cultured kelp or juveniles of natural populations is still an open question in the highly exposed Sonoma and Mendocino County coastlines George et al. (2015).

In other rocky reef systems, kelp forest restoration success has been determined by kelp introduction and removal of stressors (such as active grazers) (Morris et al., 2020). Kelp introduction was a determinant for the success of *Lessonia nigrescens* restoration in the northern coast of Chile (Correa et al., 2006). Previous work has shown that younger kelp sporophytes are more prone to predation (Lubchenco, 1983). Thus, identifying how to minimize urchin predation on younger transplants or outplants is key to ensuring successful introduction of kelp. Previous studies have proposed using grazer exclusion devices (Carney et al., 2005) or choosing sites where grazers are not as highly abundant can benefit kelp introduction (Duggins et al., 2001). The sensitivity of our model to urchin grazing rate further supports the potential efficacy of such approaches.

While our model can provide qualitative management-relevant insights into the relative efficacy of different approaches (e.g. initial vs. ongoing interventions; urchin removal, kelp reseeding, and kelp outplanting separately or in combination), quantitatively precise insights, such as the exact urchin threshold and kelp densities that can enable recovery, are more challenging due to data limitations. Our best-fit model has wide-ranging posterior distributions for most parameters (Figure B.1), including those that strongly influence the threshold urchin and kelp densities for recovery such as the urchin grazing rate on kelp. In addition, these parameters will inevitably vary in space and time, such as through urchin grazing dependence on water temperature and sedimentation (Traiger, 2019), such that no one target value will apply. Our sensitivity analysis can inform data collection aimed at resolving parameters (and their environmental dependencies) most likely to improve the ability to precisely estimate target values for urchin removal and kelp reintroduction. As more data becomes available from both kelp recovery monitoring and monitoring of potential abiotic drivers at finer spatial and temporal scales, our model (with extensions to address the assumptions described in the “Model limitations” section below) provides a foundational quantitative framework for leveraging those data for more precise predictions of threshold values for achieving a target recovery likelihood or rate.

Another consideration for kelp restoration management, especially in northern California, is the reintroduction of predators such as the sunflower seastar. A role for predator reintroduction is evident in the high sensitivity of the recovery threshold and rate in our model to parameters that likely depend on predator presence: urchin mortality (*δ_A_*) and the potential for urchins to switch between active and passive grazing (*σ_A_*). If the urchin grazing mode depends on a cryptic behavioral response to predator presence (Cowen, 1983; Duggins, 1983) as well as drift kelp presence (Harrold and Reed, 1985), then predator reintroduction could increase system resilience in terms of both likelihood and rate of recovery. Empirically, predator decline was one of the identified drivers of kelp forest loss in northern California (Rogers-Bennett and Catton, 2019; McPherson et al., 2021), such that predator reintroduction is then a component of addressing the drivers of system degradation, which is a key determinant of restoration success (Palmer et al., 2010; Morris et al., 2020). As noted in the Methods: Model overview, we did not explicitly include predator reintroduction because of uncertainty in its near-term feasibility and because initial restoration to a kelp-dominated system with non-barren, nutritious urchins through the interventions modeled here might determine the potential efficacy and optimal timing of predator reintroduction. Therefore, while our model can provide insight into restoration management on the short (annual) time scales modelled here, understanding the potential long-term recovery of ecosystem structure and resilience will likely require consideration of predator dynamics and reintroduction.

Our findings provide a system-specific case study of threshold-based approaches illuminated in previous theoretical models that look at the optimal restoration strategy of partial restoration efforts. Lampert and Hastings (2014) suggest that an economically optimal approach is to engage in restoration efforts until the target population reaches a certain threshold (the urchin threshold density in our case). In our work we find two key restoration strategies to perform at early stages of restoration: urchin removal and kelp outplanting. Lampert and Hastings (2019) further suggest that the optimal restoration strategy is to implement one strategy until a certain threshold is reached (removing urchins below the urchin threshold density), and then combining the two strategies until a certain “investment benchmark” is achieved, after which the system (e.g. kelp forest) will recover from natural processes. This benchmark might be determined in terms of a minimum kelp density or a maximum urchin density in the restored kelp oasis. Budget limitations may restrict the number of target sites that can be successfully restored (Wilson et al., 2011), which makes choosing priority sites based on likelihood of restoration success an important step when performing restoration. Finally, the three strategies explored in this work might have an optimal timing of when to be applied, which likely differs for each strategy (Lampert and Liebhold, 2021). Finding the optimal timing of application of the three strategies explored in this work and other unexplored strategies is still an open question.

### 4.2 Model limitations

As with any model, we made a number of assumptions to construct the simplest possible model relevant to our central questions. We have chosen a Laplacian dispersal kernel, but dispersal of kelp seeds is known to be highly dependent on currents, which may skew the direction of dispersal (Gaylord et al., 2004). With advection, kelp spread rate would likely increase in favor of the direction of the current (Lou and Lutscher, 2014), dependent on physical factors such as seed buoyancy and water turbulence. These physical factors could be further explored using a non-parametric kernel (Richardson et al., 2018). We also ignore adult urchin movement, where the effect will depend on urchin movement responses to kelp recovery, where urchins exhibit lower movement inside than outside kelp forests due to differences in food availability (Mattison et al., 1977). If kelp recovery reduces urchin movement due to increased drift kelp availability, then accounting for urchin movement might decrease the amount of urchin removal and the role of ongoing restoration in restoration efficacy. Alternatively, if recovering kelp attracts high-movement barren urchins as active grazers, then accounting for urchin movement might increase the amount of urchin removal and the role of ongoing restoration in restoration efficacy.

Other physical factors that are not considered explicitly in this model are variations in environmental conditions such as temperature and nutrients, which are known to affect kelp productivity and growth (Bell et al., 2015). Our sensitivity analyses show that parameters highly dependent on environmental conditions in our model such as kelp mortality (*δ_A_*) are influential in determining both the urchin density threshold and kelp recovery rate. Thus, both urchin threshold density and kelp recovery rate might be higher at regions of the coastline with environmental conditions that further enhance kelp survival, leading to location-specific restoration intervention intensity required for success. If stable in time, local variation in environmental conditions could also help identify regions of the coastline with a higher potential to become kelp oases (Heinrichs et al., 2016).

Our model also assumes that the effects of ongoing restoration effort occur instantaneously. In reality, restoration efforts might present lags in their impact for reasons such as long life-cycles of the target population (Uezu and Metzger, 2016) or natural lags in the biogeochemical cycles (Hamilton, 2012). For kelp, a lag between seed outplanting and sporophyte establishment and maturation, during when kelp outplants might be more vulnerable to urchin grazing (Anderson et al., 1997), captured in our model with a separate grazing rate *γ_S_*, could decrease the efficacy of seed outplanting modelled here.

In addition, we focus our model on the dynamics of kelp sporophytes and implicitly considers the dynamics of the gametophytes. Gametophytes have the potential to act as a spore bank similar to a terrestrial seed bank, with persistence in a dormant state for an extended period of time until favorable environmental conditions occur (Edwards, 2000). This could enable kelp recovery in the absence of the extant “kelp oases” modeled here and lead to the alternative stable states observed in our model to behave as long transients instead (Arroyo-Esquivel et al., 2022). In this case, management might focus on how to decrease the length of the urchin barren transient state while also increasing the length of the kelp forest transient state, if a goal is to avoid extended periods of economic loss from kelp-associated livelihoods such as the red urchin commercial fishery, red abalone recreational fishery, and recreational diving. While analyzing the effect of different restoration interventions on transient duration given a kelp spore bank would require model modifications, given the importance of the grazing interaction between kelp and urchins found here, we suspect the qualitative results of our model would not change significantly.

Our best-fit model also assumes that alternative stable states are relevant to the kelp-urchin dynamics observed in our system. Preliminary evidence suggests kelp is returning to some areas of the coastline, potentially due to more nutrient-rich, colder waters (B. Hughes, personal communication). This recovery could be due to a shift in environmental conditions from a range where alternative stable states were relevant to a range where the kelp-dominated is the only relevant state. Alternatively, this recovery might indicate that the system state shifts with environmental conditions without alternative stable states. While these alternative explanations affect the relevance of our results concerning a threshold value of urchin density or kelp reintroduction that enables recovery, in either case restoration might still affect the rate of recovery, as observed in northern California (Ward et al., 2022), especially if transients are slow as noted above. Determining which case best explains this apparent recovery will require analyzing the emerging data in the coming years in comparison to model predictions with environmental drivers and different model structures with or without alternative stable states (as done for giant kelp in southern California in Karatayev et al. (2021), where there is greater data availability than for the northern California bull kelp system that is our focus here).

Our model considers only the interactions between sea urchins and kelp, which are central to the efficacy of current restoration interventions. In reality, an array of other species in California’s temperate rocky reefs might affect recovery dynamics and restoration outcomes. For example, crustose coralline algae competing with kelp may also facilitate urchin recruitment, potentially decreasing the threshold urchin density, i.e. increase the urchin removal necessary for kelp recovery (Baskett and Salomon, 2010). In comparison, the presence of a natural predator of sea urchins such as the sunflower sea star (currently functionally extinct on the California north coast (Rogers-Bennett and Catton, 2019)) would lead to a more cryptic behavior of urchins, which would lead to an increase in threshold urchin density (Smith et al., 2021). Adding such predators and their potential reintroductioncan add another ecological ecological dimension to restoration, as discussed above in the section on Model implications. Therefore, further adding ecological realism can allow extensions in the capacity of our model to inform restoration efforts.

## Acknowledgements

This research is supported under California Sea Grant #R/HCE-15: “A multi-pronged approach to kelp recovery along California’s north coast”. In addition, we thank Dr. Rietta Hohman and Dr. Norah Eddy for their suggestions on restoration strategies. The authors would also like to thank three anonymous reviewers for their feedback on an earlier version of this manuscript. JAE thanks the University of Costa Rica for their support through the development of this paper.

## S1 Presence of alternate stable states in the nonspatial version of the model

In this appendix we show the presence of alternate stable states in our model by showing it has trajectories at where kelp recovers (kelp forest state) and where kelp collapses (urchin barren) at different initial conditions. To show these trajectories, we run our spatial model 1 and then calculate the total kelp density (denoted by **A**_*t*_) and urchin density (denoted by **U**_*t*_) by integrating their densities at each point *y* over the entire coastline. In mathematical terms, we calculate these total densities using the expressions:

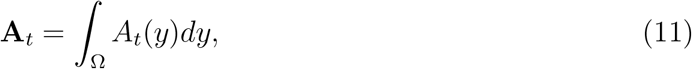

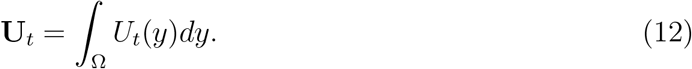

Figure 1 provides trajectories of total kelp and urchin densities after 24 time steps under a range of values for initial kelp and urchin densities (and with the same set of paramter values for all simulations). These simulations demonstrate the existence of two alterative steady-state outcomes in the model: the time series converge to either (a) a limit cycle of high kelp density and urchin persistence (kelp forest state) or (b) a barren state where kelp declines to regional extinction and the urchins initially increase but eventually collapse due to starvation. Which of these states is the long-term outcome depends on initial kelp and urchin densities relative to the threhold values identified in Fig. 3.

## S2 Posterior distributions of parameters

**Figure S 1:**
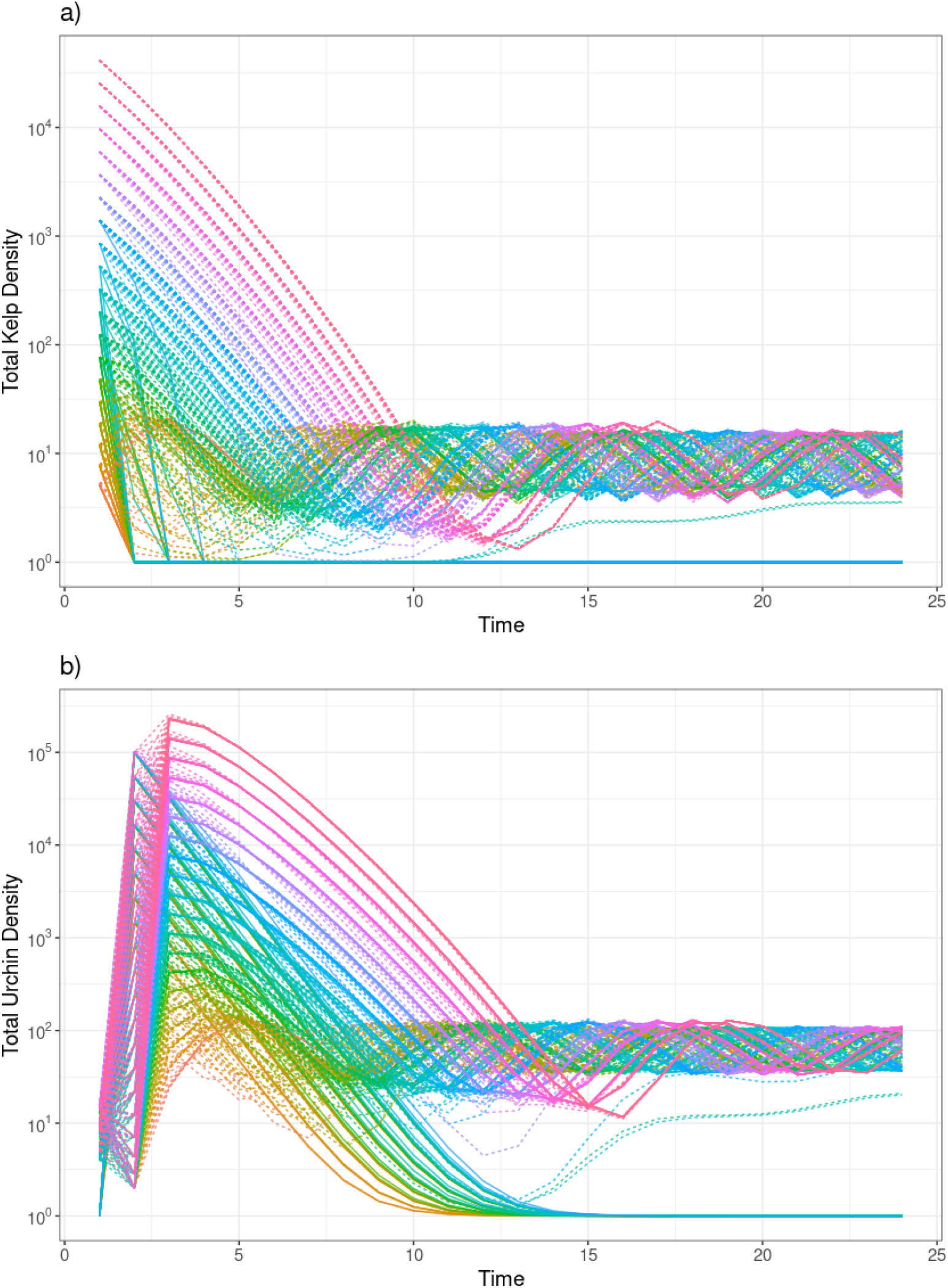
Trajectories of **a)** total kelp (**A**_*t*_) and **b)** urchin (**U**_*t*_) densities. Each color represents different initial conditions, the solid lines represent trajectories that converge into an urchin barren, while the dashed lines represent trajectories that converge into a kelp forest state.

**Figure S 2:**
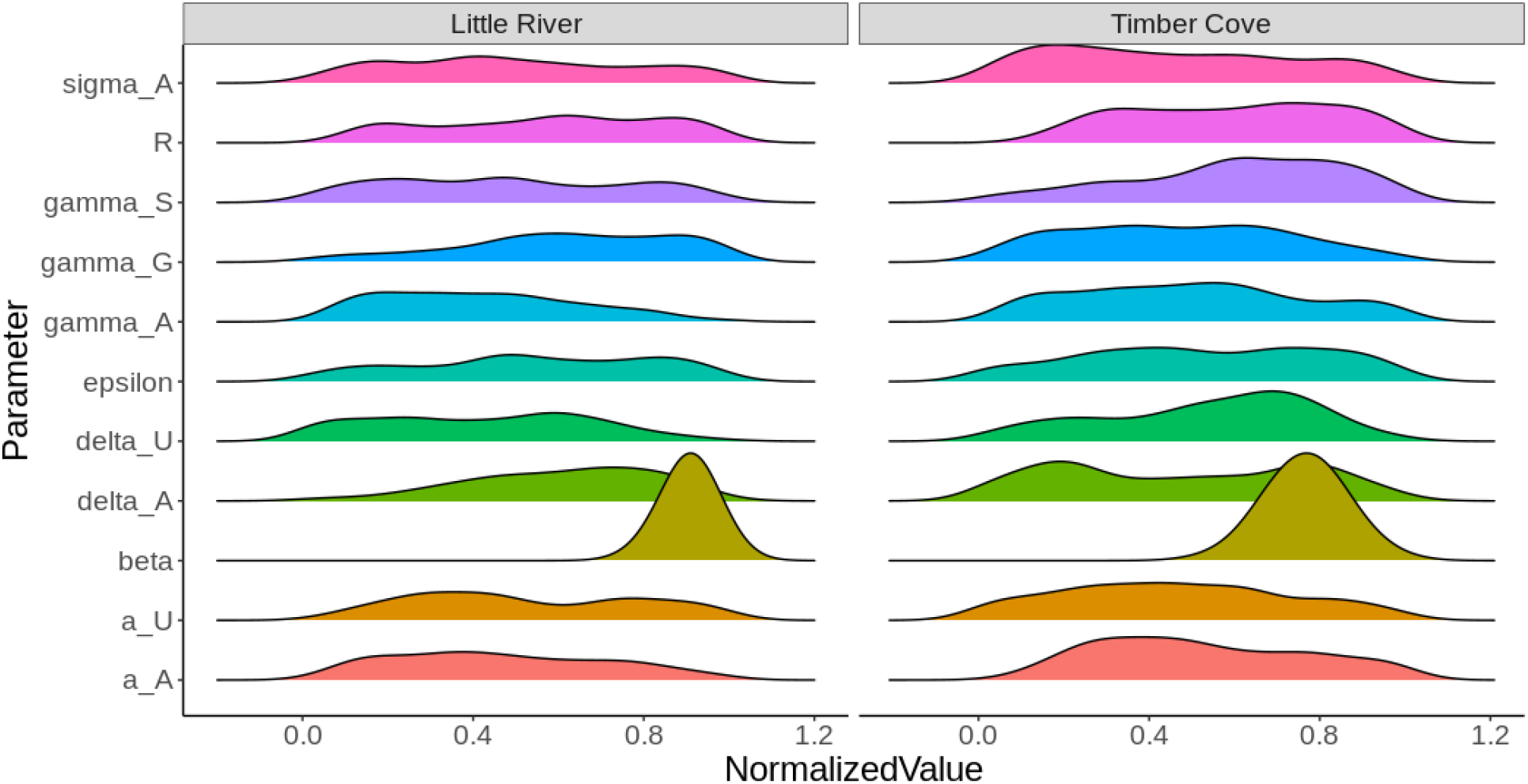
Posterior distributions obtained using the Approximate Bayesian Computation algorithm for each of the parameters for both the Little River and Timber Cove regions. Notice that *β* is estimated using a different approach (see main text).

## S3 Spread rates estimated from the model at different values of kelp mean dispersal distance1/*a_A_*

**Figure S 3:**
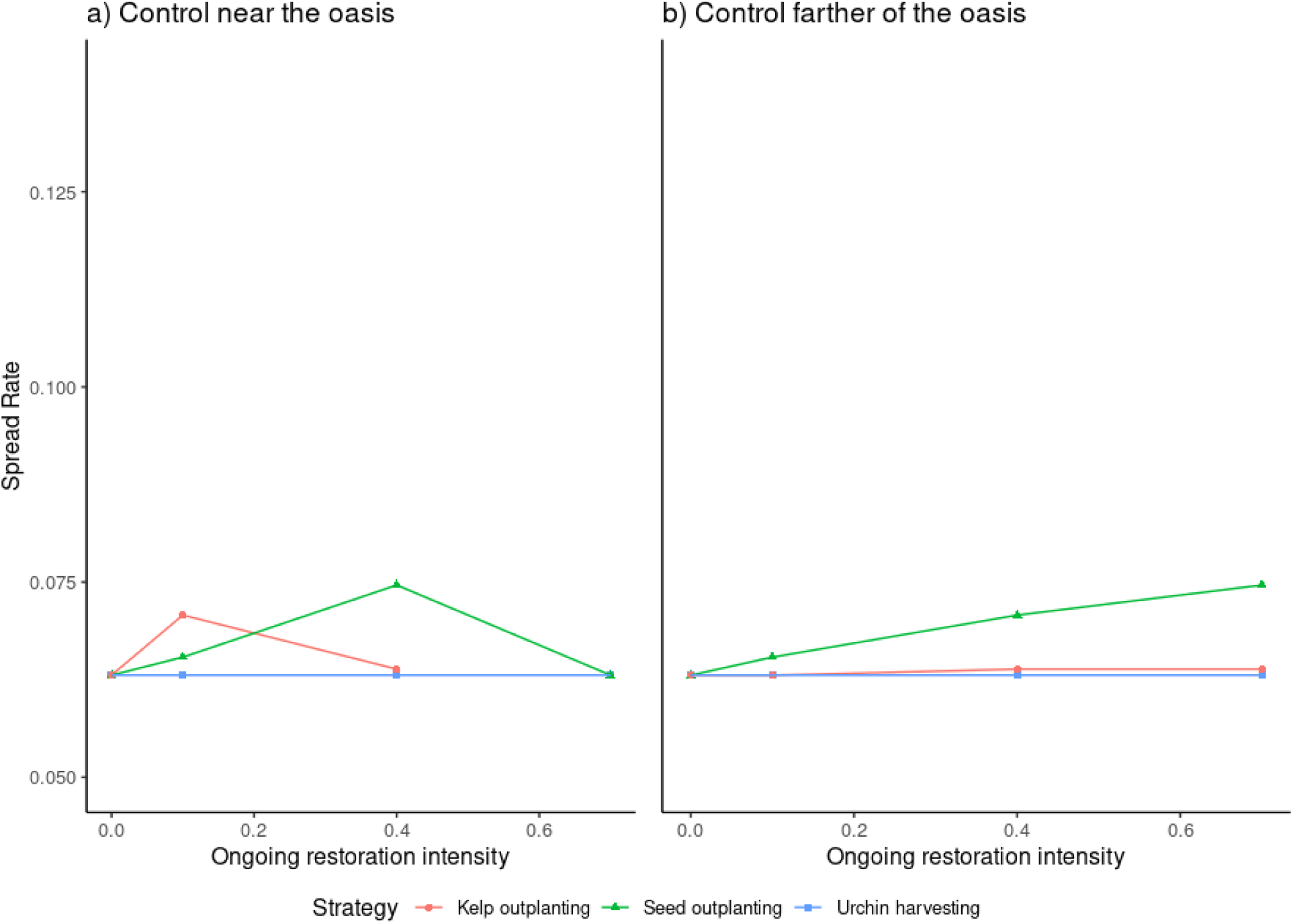
Kelp spread rate under different restoration strategies with increasing intensity following urchin removal below the threshold value necessary for kelp recovery and with an initial kelp density 2/*β* and half the estimated mean dispersal distance (double the baseline of *a_A_*). Each line represents a different strategy: kelp outplanting in red circles, kelp seeding in green triangles, and sustained urchin harvest in blue squares. Panel a) shows ongoing restoration efforts near the kelp oasis (*η* = 1), and panel b) shows ongoing restoration efforts across a wider region of the coastline (*η* = 10). Compare to Fig. 6

**Figure S 4:**
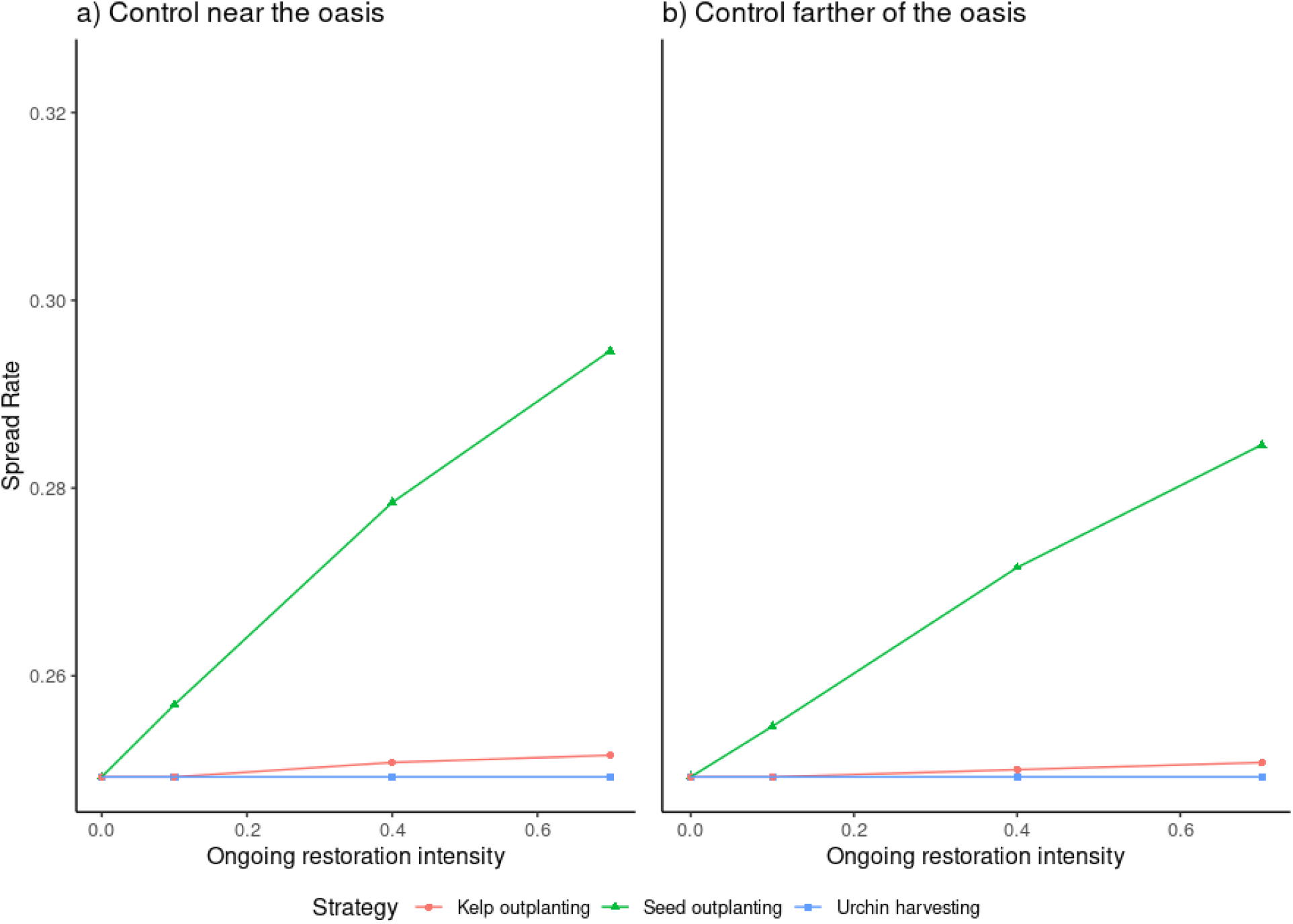
Kelp spread rate under different restoration strategies with increasing intensity following urchin removal below the threshold value necessary for kelp recovery and with an initial kelp density 2/*β* and double the estimated mean dispersal distance (half the baseline value of *a_A_*). Each line represents a different strategy: kelp outplanting in red circles, kelp seeding in green triangles, and sustained urchin harvest in blue squares. Panel a) shows ongoing restoration efforts near the kelp oasis (*η* = 1), and panel b) shows ongoing restoration efforts across a wider region of the coastline (*η* = 10). Compare to Fig. 6

**Figure S 5:**
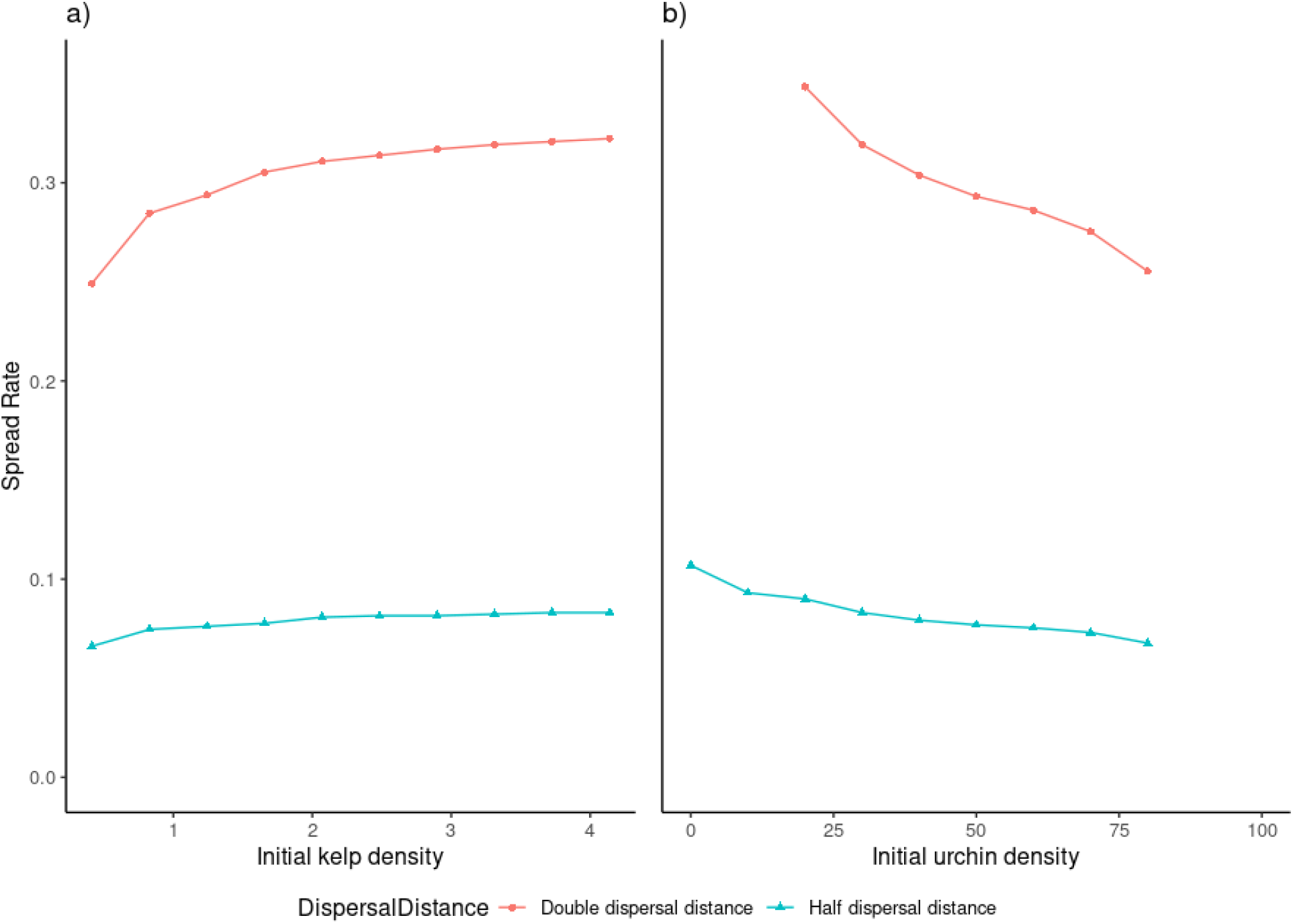
Kelp spread rate under different initial conditions for partial restoration efforts at the initial (short-term) restoration stage. Each line represents a different value of mean dispersal distance, either double or half mean dispersal distance from the baseline. Panel a) shows the change in spread rate as the initial kelp density varies, and panel b) shows the change in spread rate as the initial urchin density varies.

## References

Acosta, A. L., d’Albertas, F., Leite, M. d. S., Saraiva, A. M., and Metzger, J. P. W. (2018). Gaps and limitations in the use of restoration scenarios: a review. Restoration Ecology, 26(6):1108–1119. eprint: https://onlinelibrary.wiley.com/doi/pdf/10.1111/rec.12882.

Anderson, R. J., Carrick, P., Levitt, G. J., and Share, A. (1997). Holdfasts of adult kelp Ecklonia maxima provide refuges from grazing for recruitment of juvenile kelps. Marine Ecology Progress Series, 159:265–273.

Arroyo-Esquivel, J., Hastings, A., and Baskett, M. L. (2022). Characterizing Long Transients in Consumer–Resource Systems With Group Defense and Discrete Reproductive Pulses. Bulletin of Mathematical Biology, 84(9):102.

Baskett, M. L. and Salomon, A. K. (2010). Recruitment facilitation can drive alternative states on temperate reefs. Ecology, 91(6):1763–1773. eprint: https://esajournals.onlinelibrary.wiley.com/doi/pdf/10.1890/09-0515.1.

Beisner, B. E., Haydon, D. T., and Cuddington, K. (2003). Alternative stable states in ecology. Frontiers in Ecology and the Environment, 1(7):376–382. eprint: https://esajournals.onlinelibrary.wiley.com/doi/pdf/10.1890/1540-9295%282003%29001%5B0376%3AASSIE%5D2.0.CO%3B2.

Bell, T. W., Cavanaugh, K. C., Reed, D. C., and Siegel, D. A. (2015). Geographical variability in the controls of giant kelp biomass dynamics. Journal of Biogeography, 42(10):2010–2021. eprint: https://onlinelibrary.wiley.com/doi/pdf/10.1111/jbi.12550.

Beverton, R. J. H. and Holt, S. J. (1993). On the Dynamics of Exploited Fish Populations. Fish & Fisheries Series. Springer Netherlands.

Bowden-Kerby, A. (2001). Low-tech coral reef restoration methods modeled after natural fragmentation processes. Bulletin of Marine Science, 69(2):915–931.

Bradshaw, A. D. (1996). Underlying principles of restoration. Canadian Journal of Fisheries and Aquatic Sciences, 53(S1):3–9.

Cahall, R. E., Hayes, J. P., and Betts, M. G. (2013). Will they come? Long-term response by forest birds to experimental thinning supports the “Field of Dreams” hypothesis. Forest Ecology and Management, 304:137–149.

Carney, L., Waaland, J., Klinger, T., and Ewing, K. (2005). Restoration of the bull kelp Nereocystis luetkeana in nearshore rocky habitats. Marine Ecology Progress Series, 302:49–61.

Cochran, R. C. and Engelmann, F. (1975). Environmental regulation of the annual reproductive season of strongylocentrotus purpuratus (stimpson). The Biological Bulletin, 148(3):393–401.

Connell, S. D., Kroeker, K. J., Fabricius, K. E., Kline, D. I., and Russell, B. D. (2013). The other ocean acidification problem: CO2 as a resource among competitors for ecosystem dominance. Philosophical Transactions of the Royal Society B: Biological Sciences, 368(1627):20120442. Publisher: Royal Society.

Correa, J. A., Lagos, N. A., Medina, M. H., Castilla, J. C., Cerda, M., Ramírez, M., Martínez, E., Faugeron, S., Andrade, S., Pinto, R., and Contreras, L. (2006). Experimental transplants of the large kelp Lessonia nigrescens (Phaeophyceae) in high-energy wave exposed rocky intertidal habitats of northern Chile: Experimental, restoration and management applications. Journal of Experimental Marine Biology and Ecology, 335(1):13–18.

Cowen, R. K. (1983). The effects of sheephead (Semicossyphus pulcher) predation on red sea urchin (Strongylocentrotus franciscanus) populations: an experimental analysis. Oecologia, 58(2):249–255.

Dalrymple, S. E., Banks, E., Stewart, G. B., and Pullin, A. S. (2012). A Meta-Analysis of Threatened Plant Reintroductions from across the Globe. In Maschinski, J., Haskins, K. E., and Raven, P. H., editors, Plant Reintroduction in a Changing Climate: Promises and Perils, The Science and Practice of Ecological Restoration, pages 31–50. Island Press/Center for Resource Economics, Washington, DC.

Dobkowski, K. A., Flanagan, K. D., and Nordstrom, J. R. (2019). Factors influencing recruitment and appearance of bull kelp, *Nereocystis luetkeana* (phylum Ochrophyta). Journal of Phycology, 55(1):236–244.

Dodd, C. K. and Seigel, R. A. (1991). Relocation, Repatriation, and Translocation of Amphibians and Reptiles: Are They Conservation Strategies That Work? Herpetologica, 47(3):336–350. Publisher: [Herpetologists’ League, Allen Press].

Duggins, D., Eckman, J. E., Siddon, C. E., and Klinger, T. (2001). Interactive roles of mesograzers and current flow in survival of kelps. Marine Ecology Progress Series, 223:143–155.

Duggins, D. O. (1983). Starfish Predation and the Creation of Mosaic Patterns in a Kelp-Dominated Community. Ecology, 64(6):1610–1619.

Dumont, C. P., Himmelman, J. H., and Russell, M. P. (2006). Daily movement of the sea urchin Strongylocentrotus droebachiensis in different subtidal habitats in eastern Canada. Marine Ecology Progress Series, 317:87–99.

Ebert, T. A., Schroeter, S. C., Dixon, J. D., and Kalvass, P. (1994). Settlement patterns of red and purple sea urchins (Strongylocentrotus franciscanus and S. purpuratus) in California, USA. Marine Ecology Progress Series, 111(1/2):41–52.

Edwards, M. S. (2000). THE ROLE OF ALTERNATE LIFE-HISTORY STAGES OF A MARINE MACROALGA: A SEED BANK ANALOGUE? Ecology, 81(9):2404–2415.

Eger, A. M., Marzinelli, E., Gribben, P., Johnson, C. R., Layton, C., Steinberg, P. D., Wood, G., Silliman, B. R., and Vergés, A. (2020). Playing to the Positives: Using Synergies to Enhance Kelp Forest Restoration. Frontiers in Marine Science, 7. Publisher: Frontiers.

Folke, C., Carpenter, S., Walker, B., Scheffer, M., Elmqvist, T., Gunderson, L., and Holling, C. (2004). Regime Shifts, Resilience, and Biodiversity in Ecosystem Management. Annual Review of Ecology, Evolution, and Systematics, 35(1):557–581. eprint: https://doi.org/10.1146/annurev.ecolsys.35.021103.105711.

Ford, T. and Meux, B. (2010). Giant kelp community restoration in Santa Monica Bay. Urban Coast, 2:43–46.

Gaylord, B., Reed, D. C., Washburn, L., and Raimondi, P. T. (2004). Physical–biological coupling in spore dispersal of kelp forest macroalgae. Journal of Marine Systems, 49(1):19–39.

George, D. A., Largier, J. L., Storlazzi, C. D., and Barnard, P. L. (2015). Classification of rocky headlands in California with relevance to littoral cell boundary delineation. Marine Geology, 369:137–152.

Graham, M., Harrold, C., Lisin, S., Light, K., Watanabe, J., and Foster, M. (1997). Population dynamics of giant kelp Macrocystis pyrifera along a wave exposure gradient. Marine Ecology Progress Series, 148:269–279.

Hale, R. and Swearer, S. E. (2017). When good animals love bad restored habitats: how maladaptive habitat selection can constrain restoration. Journal of Applied Ecology, 54(5):1478–1486.

Hamilton, S. K. (2012). Biogeochemical time lags may delay responses of streams to ecological restoration: *Time lags in stream restoration*. Freshwater Biology, 57:43–57.

Harper, E. B., Stella, J. C., and Fremier, A. K. (2011). Global sensitivity analysis for complex ecological models: a case study of riparian cottonwood population dynamics. Ecological Applications, 21(4):1225–1240. eprint: https://esajournals.onlinelibrary.wiley.com/doi/pdf/10.1890/10-0506.1.

Harrold, C. and Reed, D. C. (1985). Food Availability, Sea Urchin Grazing, and Kelp Forest Community Structure. Ecology, 66(4):1160–1169. eprint: https://esajournals.onlinelibrary.wiley.com/doi/pdf/10.2307/1939168.

Harvell, C. D., Montecino-Latorre, D., Caldwell, J. M., Burt, J. M., Bosley, K., Keller, A., Heron, S. F., Salomon, A. K., Lee, L., Pontier, O., Pattengill-Semmens, C., and Gaydos, J. K. (2019). Disease epidemic and a marine heat wave are associated with the continental-scale collapse of a pivotal predator (Pycnopodia helianthoides). Science Advances, 5(1):eaau7042. Publisher: American Association for the Advancement of Science Section: Research Article.

Heinrichs, J. A., Lawler, J. J., and Schumaker, N. H. (2016). Intrinsic and extrinsic drivers of source–sink dynamics. Ecology and Evolution, 6(4):892–904.

Hohman, R., Hutto, S., Catton, C., and Koe, F. (2019). Sonoma-Mendocino Bull Kelp Recovery Plan. Plan for the Greater Farallones National Marine Sanctuary and the California Department of Fish and Wildlife.

Jabot, F., Faure, T., and Dumoulin, N. (2013). EasyABC: performing efficient approximate Bayesian computation sampling schemes using R. Methods in Ecology and Evolution, 4(7):684–687. eprint: https://besjournals.onlinelibrary.wiley.com/doi/pdf/10.1111/2041-210X.12050.

James, J. J., Sheley, R. L., Erickson, T., Rollins, K. S., Taylor, M. H., and Dixon, K. W. (2013). A systems approach to restoring degraded drylands. Journal of Applied Ecology, 50(3):730–739. eprint: https://besjournals.onlinelibrary.wiley.com/doi/pdf/10.1111/1365-2664.12090.

Johnson, C. R., Chabot, R. H., Marzloff, M. P., and Wotherspoon, S. (2017). Knowing when (not) to attempt ecological restoration. Restoration Ecology, 25(1):140–147.

Kanary, L., Musgrave, J., Tyson, R. C., Locke, A., and Lutscher, F. (2014). Modelling the dynamics of invasion and control of competing green crab genotypes. Theoretical Ecology, 7(4):391–406.

Karatayev, V. A., Baskett, M. L., Kushner, D. J., Shears, N. T., Caselle, J. E., and Boettiger, C. (2021). Grazer behavior can regulate large-scale patterns of community states. Ecology Letters, 24(9):1917–1929.

Katwijk, M. M., Thorhaug, A., Marbà, N., Orth, R. J., Duarte, C. M., Kendrick, G. A., Althuizen, I. H. J., Balestri, E., Bernard, G., Cambridge, M. L., Cunha, A., Durance, C., Giesen, W., Han, Q., Hosokawa, S., Kiswara, W., Komatsu, T., Lardicci, C., Lee, K., Meinesz, A., Nakaoka, M., O’Brien, K. R., Paling, E. I., Pickerell, C., Ransijn, A. M. A., and Verduin, J. J. (2016). Global analysis of seagrass restoration: the importance of large-scale planting. Journal of Applied Ecology, 53(2):567–578.

Krumhansl, K. A., Okamoto, D. K., Rassweiler, A., Novak, M., Bolton, J. J., Cavanaugh, K. C., Connell, S. D., Johnson, C. R., Konar, B., Ling, S. D., Micheli, F., Norderhaug, K. M., Pérez-Matus, A., Sousa-Pinto, I., Reed, D. C., Salomon, A. K., Shears, N. T., Wernberg, T., Anderson, R. J., Barrett, N. S., Buschmann, A. H., Carr, M. H., Caselle, J. E., Derrien-Courtel, S., Edgar, G. J., Edwards, M., Estes, J. A., Goodwin, C., Kenner, M. C., Kushner, D. J., Moy, F. E., Nunn, J., Steneck, R. S., Vásquez, J., Watson, J., Witman, J. D., and Byrnes, J. E. K. (2016). Global patterns of kelp forest change over the past half-century. Proceedings of the National Academy of Sciences, 113(48):13785–13790.

Lampert, A. and Hastings, A. (2014). Optimal control of population recovery – the role of economic restoration threshold. Ecology Letters, 17(1):28–35.

Lampert, A. and Hastings, A. (2019). How to combine two methods to restore populations cost effectively. Ecosphere, 10(1):e02552.

Lampert, A. and Liebhold, A. M. (2021). Combining multiple tactics over time for cost-effective eradication of invading insect populations. Ecology Letters, 24(2):279–287.

Largier, J. L. (2003). Considerations In Estimating Larval Dispersal Distances From Oceanographic Data. Ecological Applications, 13(sp1):71–89.

Layton, C., Coleman, M. A., Marzinelli, E. M., Steinberg, P. D., Swearer, S. E., Vergés, A., Wernberg, T., and Johnson, C. R. (2020). Kelp Forest Restoration in Australia. Frontiers in Marine Science, 7. Publisher: Frontiers.

Lee, Y.-F., Kuo, Y.-M., Lin, Y.-H., Chu, W.-C., Wang, H.-H., and Wu, S.-H. (2006). Composition, Diversity, and Spatial Relationships of Anurans Following Wetland Restoration in a Managed Tropical Forest. Zoological Science, 23(10):883–891.

Leinaas, H. P. and Christie, H. (1996). Effects of removing sea urchins (Strongylocentrotus droebachiensis): Stability of the barren state and succession of kelp forest recovery in the east Atlantic. Oecologia, 105(4):524–536.

Liaw, A. and Wiener, M. (2002). Classification and Regression by randomForest. 2:6.

Ling, S. D. (2008). Range expansion of a habitat-modifying species leads to loss of taxonomic diversity: a new and impoverished reef state. Oecologia, 156(4):883–894.

Ling, S. D., Scheibling, R. E., Rassweiler, A., Johnson, C. R., Shears, N., Connell, S. D., Salomon, A. K., Norderhaug, K. M., Pérez-Matus, A., Hernández, J. C., Clemente, S., Blamey, L. K., Hereu, B., Ballesteros, E., Sala, E., Garrabou, J., Cebrian, E., Zabala, M., Fujita, D., and Johnson, L. E. (2015). Global regime shift dynamics of catastrophic sea urchin overgrazing. Philosophical Transactions of the Royal Society B: Biological Sciences, 370(1659):20130269. Publisher: Royal Society.

Lockwood, D. R., Hastings, A., and Botsford, L. W. (2002). The Effects of Dispersal Patterns on Marine Reserves: Does the Tail Wag the Dog? Theoretical Population Biology, 61(3):297–309.

Lou, Y. and Lutscher, F. (2014). Evolution of dispersal in open advective environments. Journal of Mathematical Biology, 69(6-7):1319–1342.

Lubchenco, J. (1983). Littornia and Fucus: Effects of Herbivores, Substratum Heterogeneity, and Plant Escapes During Succession. Ecology, 64(5):1116–1123. eprint: https://esajournals.onlinelibrary.wiley.com/doi/pdf/10.2307/1937822.

Marzloff, M. P., Little, L. R., and Johnson, C. R. (2016). Building Resilience Against Climate-Driven Shifts in a Temperate Reef System: Staying Away from Context-Dependent Ecological Thresholds. Ecosystems, 19(1):1–15.

Mattison, J. E., Trent, J. D., Shanks, A. L., Akin, T. B., and Pearse, J. S. (1977). Movement and feeding activity of red sea urchins (Strongylocentrotus franciscanus) adjacent to a kelp forest. Marine Biology, 39(1):25–30.

Maya, P. H. M., Smit, K. P., Burt, A. J., and Frias-Torres, S. (2016). Large-scale coral reef restoration could assist natural recovery in Seychelles, Indian Ocean. Nature Conservation, 16:1–17.

McPherson, M. L., Finger, D. J. I., Houskeeper, H. F., Bell, T. W., Carr, M. H., Rogers-Bennett, L., and Kudela, R. M. (2021). Large-scale shift in the structure of a kelp forest ecosystem co-occurs with an epizootic and marine heatwave. Communications Biology, 4(1):1–9. Number: 1 Publisher: Nature Publishing Group.

Morris, R. L., Hale, R., Strain, E. M. A., Reeves, S. E., Vergés, A., Marzinelli, E. M., Layton, C., Shelamoff, V., Graham, T. D. J., Chevalier, M., and Swearer, S. E. (2020). Key Principles for Managing Recovery of Kelp Forests through Restoration. BioScience, 70(8):688–698.

Mumby, P. J., Steneck, R. S., and Hastings, A. (2013). Evidence for and against the existence of alternate attractors on coral reefs. Oikos, 122(4):481–491. eprint: https://onlinelibrary.wiley.com/doi/pdf/10.1111/j.1600-0706.2012.00262.x.

Neubert, M. G., Kot, M., and Lewis, M. A. (1995). Dispersal and Pattern Formation in a Discrete-Time Predator-Prey Model. Theoretical Population Biology, 48(1):7–43.

Okamoto, D. K., Schroeter, S. C., and Reed, D. C. (2020). Effects of ocean climate on spatiotemporal variation in sea urchin settlement and recruitment. Limnology and Oceanography, 65(9):2076–2091.

Palmer, M. A., Ambrose, R. F., and Poff, N. L. (1997). Ecological Theory and Community Restoration Ecology. Restoration Ecology, 5(4):291–300. eprint: https://onlinelibrary.wiley.com/doi/pdf/10.1046/j.1526-100X.1997.00543.x.

Palmer, M. A., Menninger, H. L., and Bernhardt, E. (2010). River restoration, habitat heterogeneity and biodiversity: a failure of theory or practice? Freshwater Biology, 55:205–222.

Petraitis, P. S. and Dudgeon, S. R. (2004). Detection of alternative stable states in marine communities. Journal of Experimental Marine Biology and Ecology, 300(1):343–371.

Ratajczak, Z., Nippert, J. B., and Ocheltree, T. W. (2014). Abrupt transition of mesic grassland to shrubland: evidence for thresholds, alternative attractors, and regime shifts. Ecology, 95(9):2633–2645. eprint: https://esajournals.onlinelibrary.wiley.com/doi/pdf/10.1890/13-1369.1.

ReefCheck (2020). Global Reef Tracker.

Reich, P. and Lake, P. S. (2015). Extreme hydrological events and the ecological restoration of flowing waters. Freshwater Biology, 60(12):2639–2652. eprint: https://onlinelibrary.wiley.com/doi/pdf/10.1111/fwb.12508.

Richardson, P. J., Lundholm, J. T., and Larson, D. W. (2010). Natural analogues of degraded ecosystems enhance conservation and reconstruction in extreme environments. Ecological Applications, 20(3):728–740. eprint: https://esajournals.onlinelibrary.wiley.com/doi/pdf/10.1890/08-1092.1.

Richardson, R., Kottas, A., and Sansó, B. (2018). Bayesian non-parametric modeling for integro-difference equations. Statistics and Computing, 28(1):87–101.

Rogers-Bennett, L. and Catton, C. A. (2019). Marine heat wave and multiple stressors tip bull kelp forest to sea urchin barrens. Scientific Reports, 9(1):15050. Number: 1 Publisher: Nature Publishing Group.

Sanderson, J. C. (2003). Restoration of String Kelp (Macrocystispyrifera) habitat on Tasmania’s east and south coasts.Final report to NHT for Seacare. Report, Seacare, Hobart.

Selkoe, K. A., Blenckner, T., Caldwell, M. R., Crowder, L. B., Erickson, A. L., Essington, T. E., Estes, J. A., Fujita, R. M., Halpern, B. S., Hunsicker, M. E., Kappel, C. V., Kelly, R. P., Kittinger, J. N., Levin, P. S., Lynham, J. M., Mach, M. E., Martone, R. G., Mease, L. A., Salomon, A. K., Samhouri, J. F., Scarborough, C., Stier, A. C., White, C., and Zedler, J. (2015). Principles for managing marine ecosystems prone to tipping points. Ecosystem Health and Sustainability, 1(5):1–18. Publisher: Taylor & Francis eprint: https://doi.org/10.1890/EHS14-0024.1.

Severns, P. M. (2011). Habitat restoration facilitates an ecological trap for a locally rare, wetland-restricted butterfly. Insect Conservation and Diversity, 4(3):184–191.

Smith, J. G., Tomoleoni, J., Staedler, M., Lyon, S., Fujii, J., and Tinker, M. T. (2021). Behavioral responses across a mosaic of ecosystem states restructure a sea otter–urchin trophic cascade. Proceedings of the National Academy of Sciences, 118(11). Publisher: National Academy of Sciences Section: Biological Sciences.

Springer, Y. P., Hays, C. G., Carr, M. H., and Mackey, M. R. (2010). Toward Ecosystem-Based Management of Marine Macroalgae—The Bull Kelp, Nereocystis Luetkeana. In Oceanography and Marine Biology.

Stoddard, M. A., Miller, D. L., Thetford, M., and Branch, L. C. (2019). If you build it, will they come? Use of restored dunes by beach mice. Restoration Ecology, 27(3):531–537. eprint: https://onlinelibrary.wiley.com/doi/pdf/10.1111/rec.12892.

Suding, K. N. and Hobbs, R. J. (2009). Threshold models in restoration and conservation: a developing framework. Trends in Ecology & Evolution, 24(5):271–279.

Tockner, K., Schiemer, F., and Ward, J. V. (1998). Conservation by restoration: the management concept for a ri7er-floodplain system on the Danube Ri7er in Austria. page 16.

Traiger, S. B. (2019). Effects of elevated temperature and sedimentation on grazing rates of the green sea urchin: implications for kelp forests exposed to increased sedimentation with climate change. Helgoland Marine Research, 73(1):5.

Trowbridge, W. B. (2007). The Role of Stochasticity and Priority Effects in Floodplain Restoration. Ecological Applications, 17(5):1312–1324. eprint: https://esajournals.onlinelibrary.wiley.com/doi/pdf/10.1890/06-1242.1.

Uezu, A. and Metzger, J. P. (2016). Time-Lag in Responses of Birds to Atlantic Forest Fragmentation: Restoration Opportunity and Urgency. PLOS ONE, 11(1):e0147909.

Ward, M., McHugh, T., Elsmore, K., Esgro, M., Ray, J., Murphy-Cannella, M., Norton, I., and Freiwald, J. (2022). Restoration of North Coast Bull Kelp Forests: A Partnership Based Approach. Reef Check Foundation, Marina del Rey, CA.

Watanuki, A., Aota, T., Otsuka, E., Kawai, T., Kuwahara, H., and Fujita, D. (2010). Restoration of kelp beds on an urchin barren : Removal of sea urchins by citizen divers in southwestern Hokkaido. Bulletin of the Fisheries Research Agency, 32:83–87.

Wegmann, D., Leuenberger, C., and Excoffier, L. (2009). Efficient Approximate Bayesian Computation Coupled With Markov Chain Monte Carlo Without Likelihood. Genetics, 182(4):1207–1218.

Wiens, J. (1992). What Is Landscape Ecology, Really. Landscape Ecology, 7(3):149–150. Place: Lelystad Publisher: S P B Academic Publishing Bv WOS:A1992JW40100001.

Williams, J. P., Claisse, J. T., Ii, D. J. P., Williams, C. M., Robart, M. J., Scholz, Z., Jaco, E. M., Ford, T., Burdick, H., and Witting, D. (2021). Sea urchin mass mortality rapidly restores kelp forest communities. Marine Ecology Progress Series, 664:117–131.

Wilson, K. A., Lulow, M., Burger, J., Fang, Y.-C., Andersen, C., Olson, D., O’Connell, M., and McBride, M. F. (2011). Optimal restoration: accounting for space, time and uncertainty. Journal of Applied Ecology, 48(3):715–725.

Wodika, B. R. and Baer, S. G. (2015). If we build it, will they colonize? A test of the field of dreams paradigm with soil macroinvertebrate communities. Applied Soil Ecology, 91:80–89.

